# Effects of elevation and microclimatic temperatures on butterfly-flower interaction networks in a Mediterranean mountain range

**DOI:** 10.1101/2025.01.10.632329

**Authors:** Mario Alamo, Hugo Alejandro Álvarez, Mario Mingarro, Robert J. Wilson

## Abstract

1. Researching the properties of mutualistic networks over environmental gradients is a promising but underexplored means to test how global change can affect ecosystem assembly and functioning. We examined how elevation and microclimate influenced butterfly-flower interaction networks at the hottest time of year in a Mediterranean mountain range.
2. Throughout July 2023, we recorded weekly butterfly-flower interaction networks from 36 transects in nine sites, across an 800 m elevation gradient in the Sierra de Guadarrama (Central Spain). We quantified the connectance, nestedness, modularity and robustness to species loss of networks, and related these descriptors using Generalized Additive Mixed Models to elevation, microclimate temperature (modelled using Microclima), date and time of day.
3. The networks were dominated at all sites by one or two abundant butterfly and flower species, but these varied with elevation. Butterfly networks were more robust to plant species loss at higher elevations, where communities showed increased linkage density. Butterfly-plant networks became less nested at higher microclimatic temperatures in July. Network properties also varied through the day, with connectance decreasing markedly from morning to afternoon.
4. In the Mediterranean mountains studied, summer butterfly-flower interaction networks were more resilient to disturbance at high elevations and in cooler microclimates. Nectar availability could become an important limiting factor for insects in a warming climate, and understanding the mechanisms influencing the properties of flower visitor networks is therefore likely to become increasingly important for adapting the conservation of insects to climate change.

## INTRODUCTION

Species interaction networks are a valuable but under-used tool to assess impacts of climate change on ecosystems (Bascompte, 2009; Bellard et al., 2012). Ecological networks are shaped by the intrinsic properties of constituent species and their interactions (Soares et al., 2017), but also by ecological and environmental filters (Tylianakis et al., 2008; Luna et al., 2023). How these factors influence network structure, function and dynamics could help to determine how global change may disrupt ecosystems (Thébaut & Fontaine, 2010; Hong et al., 2022). However, a lack of evidence regarding the factors influencing ecological network assembly limits our ability to predict network responses to climate change (Losapio et al., 2015; Valdovinos, 2019).

Mutualistic networks vary in complexity, including differences in nestedness and compartmentalization (Bascompte, 2009), influencing their robustness to environmental perturbations (Fortuna & Bascompte, 2006). In nested networks, specialists interact with generalists: generalist species with broad abiotic and biotic niches, that interact with many other species, confer stability against random species loss (Bastolla et al., 2009; Thébault & Fontaine, 2010; Galiana et al., 2023). In addition, structural compartmentalization or modularity can limit the spread of environmental perturbations across an ecosystem (Montoya et al., 2006). Understanding how these intrinsic properties respond to abiotic changes over environmental gradients can help elucidate the factors influencing network sensitivity to change (Pellissier et al., 2018).

Elevation gradients represent a specific case in which space-for-time substitutions of ecological responses can be used to infer possible effects of climate change (Hodkinson, 2005; Sundqvist et al., 2013). For insects and plants, increasing elevation leads to more adverse conditions (e.g., shorter growing seasons, extreme winter temperatures, or extreme seasonality), commonly leading to reductions in species richness and abundance (McCain & Grytnes, 2010), and hence reduced competition, more generalist use of resources and less structured networks (Lara-Romero et al., 2019). Mountain gradients have been used to show the influence of local temperature and extreme climate events on pollinator network specialization and robustness against extinctions (Hoiss et al., 2015; Classen et al., 2020; Luna et al., 2023). Over elevation gradients, seasonal climatic variation is an important factor influencing insect-plant interaction networks (Cuartas-Hernández & Medel, 2015; Dzekashu et al., 2023), because the interactions vary in response to the phenology and abundance of insects and flowers (CaraDonna et al., 2017). Plant-pollinator networks also differ at a local scale in response to the wide variation in local climate and habitat provided by mountains (Adedoja et al., 2018; Wang et al., 2024). In this context, there is a need to determine the capacity for local climatic variation to mitigate exposure to climatic extremes and foster the resilience of species interaction networks (Burkle & Alarcón, 2011; Bhandary et al., 2023; Scheffers et al., 2014).

In this study, we test how the structure of a plant-pollinator interaction network varies over a mountain elevation gradient, in order to understand the factors influencing network robustness to climate change. We focused on interactions between butterflies and flowers in a Mediterranean mountain range at the time of year when conditions become increasingly hot and dry (July) immediately after an annual peak in butterfly diversity and abundance (Stewart et al., 2020). July is a sensitive phenological point when both the availability of pollinating insects begins to decrease, and when nectar availability for water and energy limits insect activity (Lebeau et al., 2016a, b). We quantify metrics related to the resilience of butterfly-flower interaction networks, including connectance, linkage density, vulnerability, generality, nestedness, modularity, and robustness (Landi et al., 2018). Our study covers an 800 m gradient (940-1730 m.a.s.l.) in which butterfly diversity reaches a peak (Gutiérrez et al., 2016) and habitat and topographic heterogeneity provide microclimatic buffering against warming to existing communities (Álvarez et al., 2024). The overarching aim of our study is to test how the structure of summer butterfly-flower interaction networks varies both with elevation and local temperature, to understand the factors influencing network robustness to climate change.

## MATERIALS AND METHODS

### Study area and sampling

The study was conducted in the Sierra de Guadarrama, an approximately 80 km long mountain range in the Iberian Peninsula that reaches a maximum elevation of 2428 m (40°20′ N, 4°40′ W south-west, 41°28′ N, 3°36′ W north-east) (Figure 1). The Sierra de Guadarrama features an elevation gradient characterized by its environmental heterogeneity (Tellería, 2020), encompassing four of the six bioclimatic classifications of the Mediterranean region: mesomediterranean, supramediterranean (1000-1200 m), oromediterranean (1600-2000 m) and cryomediterranean (2700-3000 m) (Rivas-Martínez, 1987).

**Figure 1.**
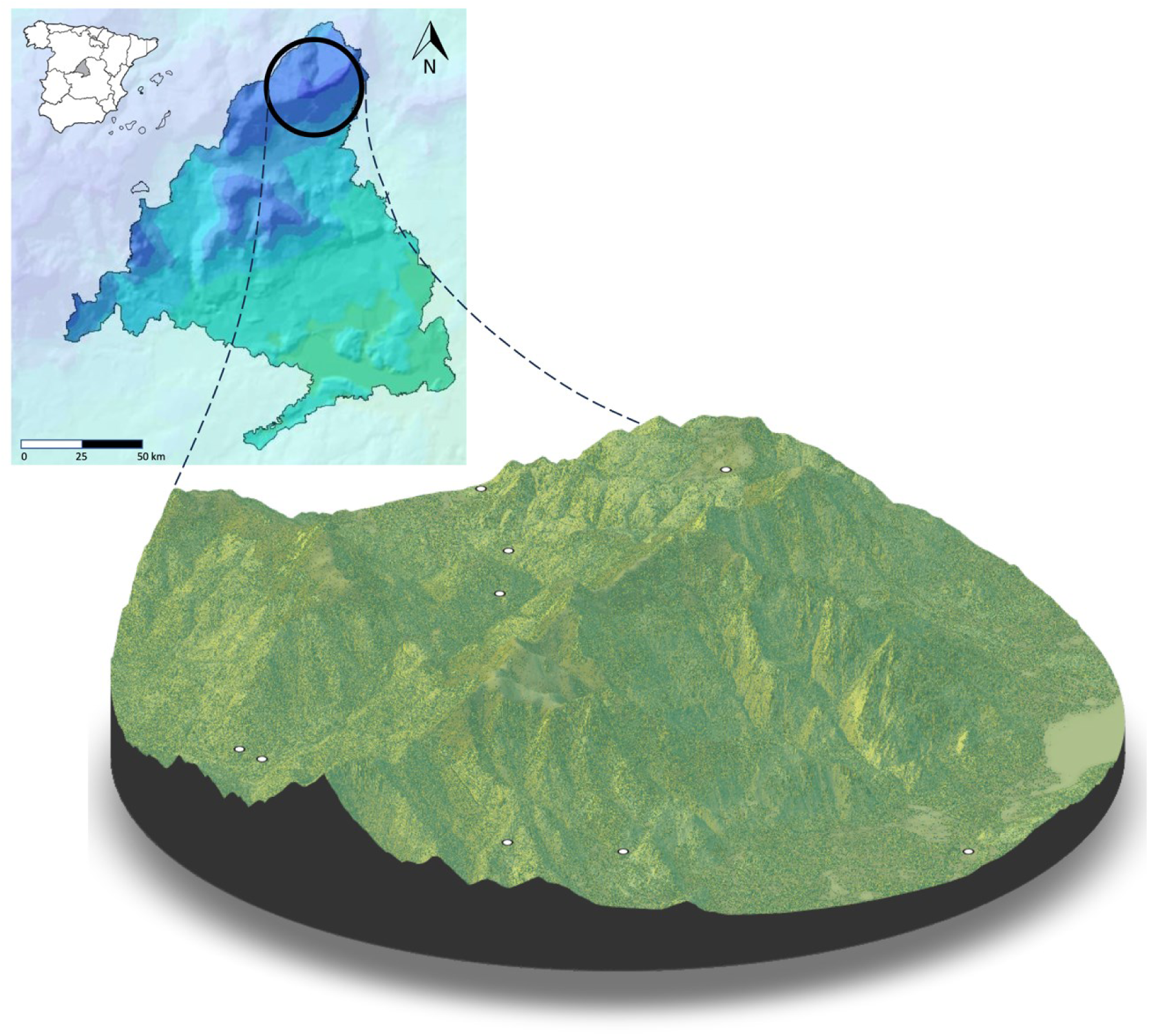
3D map of the Sierra de Guadarrama. The map shows the location of the Sierra de Guadarrama in the Madrid autonomous region, Spain. White dots represent the 9 sites where transects were set (4 transects per site) showing the elevational gradient studied in the approximately 50 km diameter region of study. The map was produced with fine resolution-LiDAR based Digital Terrain Models and Digital Vegetation Models at 5m using “rayshader” package in R software.

Sampling was conducted over four weeks in July 2023 on 36 transects across 9 sites (from highest to lowest: S1-1730 m, S2-1436 m, S3-1421m, S4-1341 m, S5-1321 m, S6-1250 m, S7-1227m, S8-1166 m and S9-941 m; Figure 1, Supplementary Table S1). Four transects (20 m long x 5 m wide, separated by at least 100 m) were established at each site within grassland clearings in forests and scrublands, spanning the oromediterranean and supramediterranean levels. These transects were selected from a wider set of butterfly communities sampled in 2004/05 and 2017 (Gutiérrez Illán et al., 2010; Álvarez et al., 2024), to provide representative coverage of environmental heterogeneity at each site and to maximize floral plant richness for observing butterfly-flower interactions. Transects were located in early July to include both open and unopened flowers, ensuring data collection throughout the four-week sampling period as conditions became hotter and drier. During the sampling period, interactions were recorded until the end of the flowering period for most available nectar plant species. Each transect was visited once a week (i.e., four consecutive visits in July, *n* = 144 replicates) during favorable conditions for butterfly activity, which included sun and low wind speed, at times between 09:00 h and 17:00 h CET (Pollard & Yates, 1993).

We recorded butterfly-flower interactions and butterfly abundance in 10-minute surveys for each transect. The observer maintained their position at the midpoint of the transect, observing all butterfly-flower interactions using Pentax Papilio II 6.5 x 21 binoculars and only moving closer to confirm butterfly identification. An interaction was recorded when a butterfly was observed feeding on a floral unit. A new interaction was considered when a butterfly interacted with a floral unit for the first time. Butterfly species were identified visually, avoiding interference with the activity of other butterflies on the transect. If necessary, butterflies were captured, identified, and released. *Pyrgus* Hübner, 1819, was the only genus that could not be identified at the species level in the field, but we kept the two observed interactions of *Pyrgus sp*. in the analysis at the genus level. To estimate overall butterfly abundance and species richness, we recorded butterflies passing through in flight during each transect sampling period. Plant identification was conducted using identification keys and distribution data (i.e., Castroviejo, 1986; Grijalbo Cervantes, 2019; Affouard et al., 2023).

### Microclimate

R packages Microclima (Maclean et al., 2019) and NicheMapR (Kearney & Porter, 2017) were used to model effects of topography and vegetation on microclimatic temperature. The two models provide a mechanistic model for the effects of vertical airflow and physical forcing on near-ground temperatures. This approach results in models that can estimate hourly temperatures with fine spatial resolution based on high temporal resolution climate data (Kearney et al., 2020), and provides accurate estimates of near-ground temperature maxima measured using dataloggers in grassland and scrubland in the Sierra de Guadarrama (Gómez-Vadillo et al., 2022). Using the runauto function of the Microclima package, we modelled temperatures for each of the 36 transects, using the centroid of each transect. Microclimate modeling was based on open scrubland "H7" (Kearney et al., 2020), an intermediate openness class for the main vegetation in our study sites. Soil and canopy albedo were set to 0.15 and 0.23 as default parameters. Hourly mean temperatures at 10 cm above the ground were estimated at a resolution of 30 m, followed by the calculation of monthly maximum daily temperatures for June (preceding sampling) and July 2023 (during sampling). Microclimatic information for June and July were selected to perform further analysis because of the importance of early summer temperatures for butterfly phenology and dynamics in the region (Gutiérrez & Wilson, 2021), and specifically for their effects on butterfly activity during our sampling period.

### Data analysis

The R package Bipartite (Dormann et al., 2008, 2009) was used to calculate seven network descriptors from quantitative bipartite interaction matrices to characterize the butterfly-flower interaction networks:

i. Connectance – the proportion of links in a network relative to the number of possible interactions. An increase in connectance indicates an increase in the generalism of the species involved (Tylianakis et al., 2010).
ii. Weighted NODF (Nestedness based on Overlap and Decreasing Fill) (Almeida-Neto & Ulrich, 2011), a descriptor evaluating the extent to which specialized species interact with subsets of the species that generalists interact with, considering the intensity of interactions between species. The weighted approach of nesting takes into account individual differences in interactions, such as preferences for specific nectar sources (Miranda et al., 2019). It is a standardized and size-invariant measure of nestedness that ranges from 1 (no nestedness) to 100 (perfect nestedness) (Ulrich et al., 2009). More nested values (higher wNODF) indicate stability and resilience to secondary extinction in mutualistic interaction networks (Tylianakis et al., 2010).
iii. Modularity (Q) refers to the degree to which interactions between different trophic levels are grouped into modules (Dormann & Strauss, 2014).
iv. Robustness is a quantitative measure of the system’s resilience to species loss. Values close to 1 indicate a system in which the elimination of species at one trophic level has minimal impact on the other levels, while values close to 0 indicate a fragile system with high dependence on species loss between trophic levels (Burgos et al., 2007).
v. Generality – the mean effective number of plant species as trophic resources available to insect species (Bersier et al., 2002).
vi. Vulnerability – the mean effective number of insect species interacting per plant species (Bersier et al., 2002).
vii. Linkage density – a quantitative measure of the diversity of interactions per species in a trophic network, calculated as the average of vulnerability and generality (Bersier et al., 2002).

Network metrics can be affected by sampling parameters, such as the size of the sampled patch and the sampling intensity (Rivera-Hutinel et al., 2012). Although our sampling design keeps these parameters constant, we used a null-model approach to test whether the observed networks differed from randomness before analyzing network metrics. The null-model approach was only applied to weighted NODF, generality, vulnerability, modularity, and robustness, as these descriptors pertain to the network structure and internal dynamics. We generated 1000 random matrices using the Patefield algorithm, which fixes the marginal sums while shuffling species interactions (Patefield, 1981). Due to the correlations of modularity and nestedness with connectance (Fortuna et al., 2010), we also employed the Vaznull algorithm, which maintains constant connectance (Vázquez et al., 2007). Observed network descriptors were compared to the random values generated by the null models using a *t*-test with the R package car (Fox & Weisberg, 2019). All observed network metrics were significantly different from the null models (Supplementary material Table S2, Table S3).

Collinearity of predictor variables is known to influence model estimation and prediction (Dormann et al., 2013). We produced pairwise Spearman Rank coefficient correlograms for all explanatory variables prior to running models using the R package GGally (Schloerke et al., 2021), to inspect collinearity based on a threshold of correlation coefficients rs>0.7 between predictor variables as an indicator. No correlation coefficients exceeded rs > 0.3 apart from July microclimate Tmax and elevation (rs = -0.6) and butterfly abundance and richness (rs = 0.7).

Generalized additive mixed models (GAMMs) were used to model the relationships between observed network metrics and environmental variables, using the R package mgcv (Wood, 2011). The explanatory variables we considered were elevation (m), mean daily maximum temperatures (°C) of modelled microclimate on each transect in June and July, date of the visit, and the time of day when sampling was undertaken. The network descriptors used as response variables were connectance (log transformed for normality); linkage density; weighted NODF; modularity Q; generality; vulnerability; and robustness on two levels: HL (butterflies) and LL (plants). To assess effects of diurnal changes in butterfly behavior on network properties, we included the variable "time of day". In addition, we tested how butterfly species richness (recorded during each transect) varied with date, time, and over the elevation gradient.

The relationships of network metrics with environmental variables were modeled using the restricted maximum likelihood method (REML) and a smoothing function with a third-order penalty (*k* = 3). By restricting the flexibility of the model fit, we balanced the modelling of the variables and the complexity of the model. Gaussian distribution was used for robustness HL, robustness LL, connectance and modularity Q. Poisson distribution was used for butterfly richness. Quasi-Poisson distribution was used for weighted NODF, linkage density, generality, and vulnerability to account for overdispersion. Finally, to control for spatial pseudo-replication, we included transect ID and site ID as a nested random effect.

All statistical analyses were performed using R software v 4.3.1 (R Core Team, 2023).

## RESULTS

### Network Composition

A total of 1145 butterfly-flower interactions were recorded. The overall network showed no pattern of compartmentation (Figure 2A) and the interactions involved: (i) 54 butterfly species from 5 families, with 87% of interactions recorded for Nymphalidae (59%) and Lycaenidae (28%); and (ii) 52 species of flowering plants from 19 families, with 82% of interactions recorded for Asteraceae (53%) and Rosaceae (29%) (Figure 2b; Supplementary Table S1). Among butterflies (first-order consumers), 40% of interactions were represented by *Melanargia lachesis* (Nymphalidae) (17%), *Lycaena virgaureae* (Lycaenidae) (13%), and *Issoria lathonia* (Nymphalidae) (10%) (Figure 2b). Among plants (producers), 57% of interactions involved *Rubus ulmifolius* (Rosaceae) (27%), *Carduus acanthoides* (Asteraceae) (17%), and *Jacobaea vulgaris* (Asteraceae) (13%) (Figure 2b).

**Figure 2.**
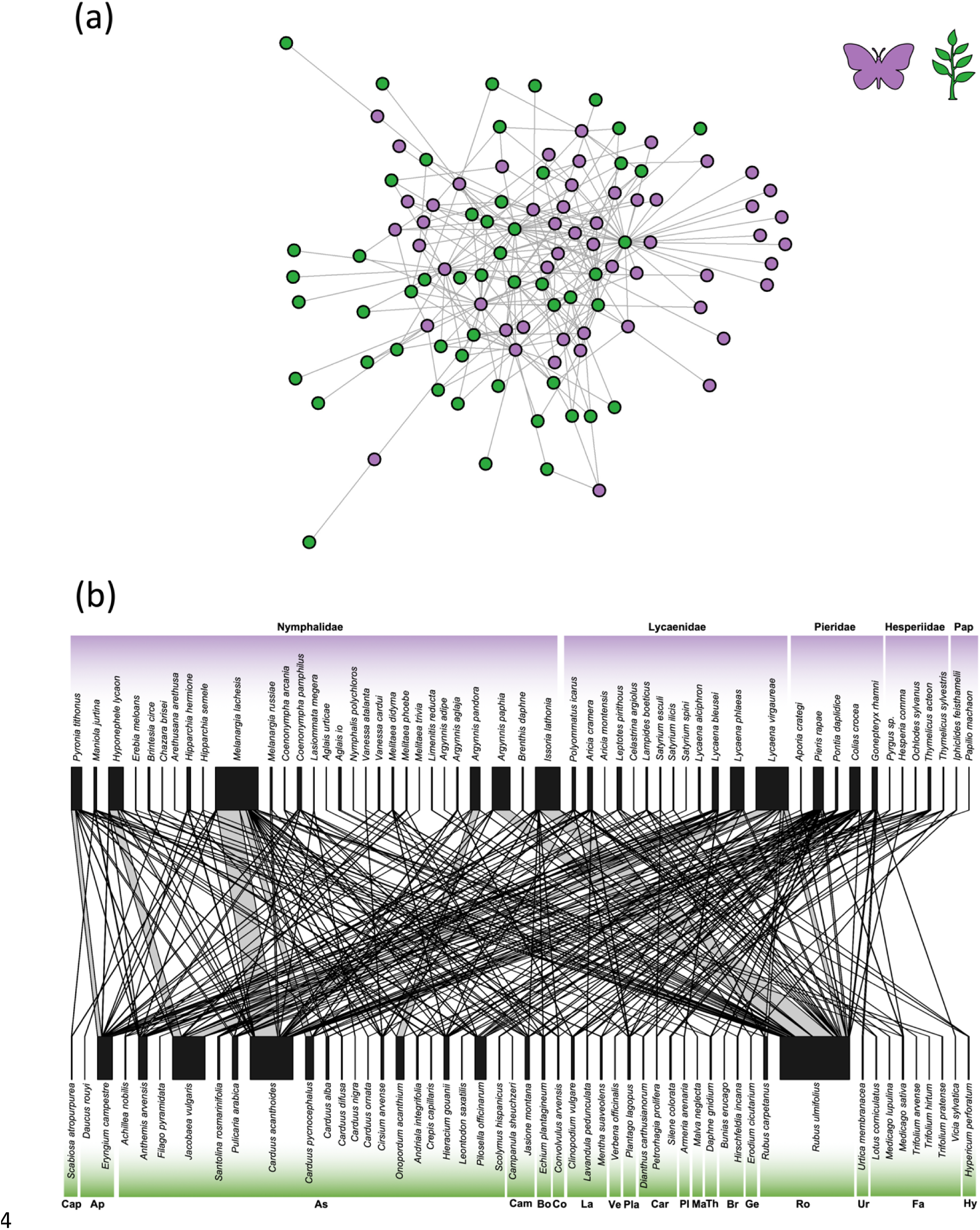
Overall quantitative interaction network, representing the level of interactions of the overall butterfly-flower trophic network in the Sierra de Guadarrama for an 800m elevational gradient. Purple represents the butterfly species (primary consumers) and green represents flower species (producers), the lines between them represent quantitative trophic interactions. Modular representation of the network showing no pattern of compartmentation (a). Bipartite representation of the network (b). The size of the lines and the nodes shows the number of interactions recorded. Species are arranged by phylogenetic relationship at the species level (butterflies) and subfamily level (plants). Butterfly families: Nymphalidae, Lycaenidae, Pieridae, Hesperiidae, and Papilionidae (Pap). Plant families: Caprifoliaceae (Cap), Apiaceae (Ap), Asteraceae (As), Campanulaceae (Cam), Boraginaceae (Bo), Convolvulaceae (Co), Lamiaceae (La), Verbenaceae (Ve), Plantaginaceae (Pla), Caryophyllaceae (Car), Plumbaginaceae (Pl), Malvaceae (Ma), Thymelaeaceae (Th), Brassicaceae (Br), Geraniaceae (Ge), Rosaceae (Ro), Urticaceae (Ur), Fabaceae (Fa) and Hypericaceae (Hy).

### Network composition at different elevations

Network size in the sampling transects (the number of butterfly and plant species recorded on each visit per transect) ranged from 3 to 18, and did not vary significantly with elevation (*n* = 144, edf = 1.524, *p* = 0.1425) (Figure 3). Most interactions at each site (composed of four sampling transects) were formed by one or two locally abundant butterfly and plant species (Supplementary Table S1), but the main interaction partners varied with elevation. The three highest sites accounted for 50% of interactions overall, with most involving butterfly species such as *L. virgaureae*, *Hyponephele lycaon* (Nymphalidae) and herbaceous plants (mainly Asteraceae) such as *J. vulgaris* and *C. acanthoides*. At intermediate elevations, the most common interactions were the nymphalid butterflies *Argynnis pandora* and *A. paphia* with *R. ulmifolius* (blackberry). At the lowest sites, the most common interactions were between *Pyronia tithonus* (Nymphalidae) and *Eryngium campestre* (Apiaceae).

**Figure 3.**
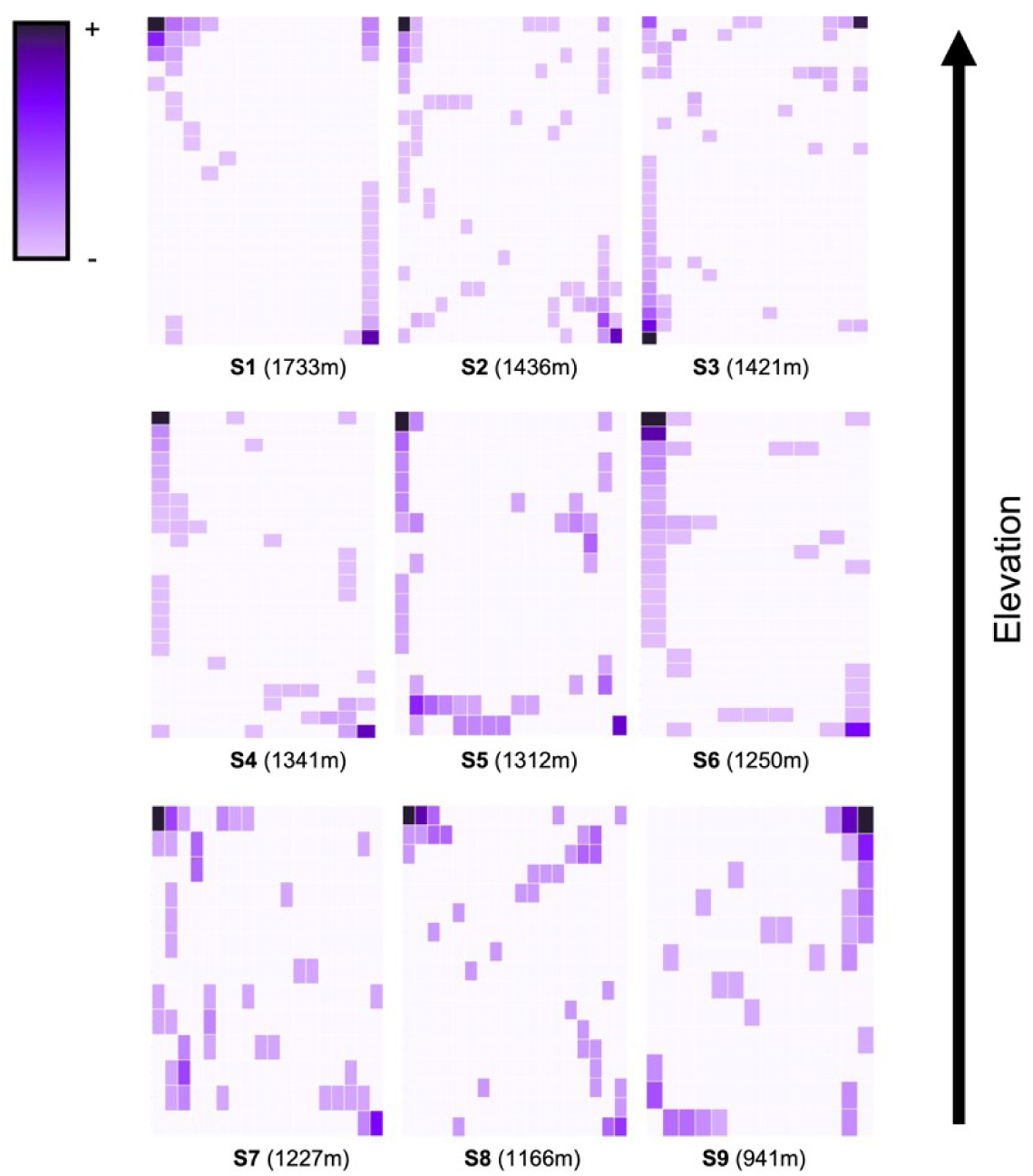
Heatmap representations of the plant-butterfly trophic network along the elevation gradient (9 sites). The top-left heatmap represents the highest elevation site and the bottom-right represents the lowest site. Columns represent plant species, and rows represent butterfly species. The intensity of butterfly-plant interactions is indicated by the color palette.

### Microclimatic Temperatures

Mean maximum daily temperatures estimated at 10 cm above the ground for the nine sites were 34°C in June and 37°C in July. July microclimatic temperature decreased by 0.4°C per 100 m increase in elevation (negative linear trend with elevation, *n* = 36 transects, *t* = -5.518, *p* < 0.001). However, June microclimatic temperatures did not show a significant trend with elevation (*n* = 36, *t* = 0.394, *p* = 0.696): this corresponds with cloudy conditions and cooler temperatures recorded in June 2023 at meteorological stations, which showed a shallower elevational trend in temperature than in July (Supplementary material Figure S1).

### Network Descriptors and Environmental Variables

In the GAMMs, four network descriptors showed significant relationships with environmental factors: robustness HL (higher level, butterflies), weighted NODF (nestedness), connectance, and linkage density (Table 1). Robustness HL had a positive relationship with elevation (Figure 4a), indicating that as elevation increased, butterfly networks exhibited greater resilience to the loss of flowering plant species. Nestedness (weighted NODF) showed a negative response to July temperature, with little nestedness at transects with temperature maxima above 37°C (Figure 4b), corresponding to six of the seven sites below 1500 m.a.s.l. (Supplementary material, Figure S1). Connectance showed a negative relationship with Time of Day (Figure 4c), with a higher proportion of possible interactions realized during the earlier hours of the day. Finally, linkage density had a positive relationship with elevation (Figure 4d), showing a greater number of butterfly-flower interactions at higher elevations. The remaining network descriptors (generality, vulnerability, robustness LL [lower level, flowering plants], and modularity) did not show significant relationships with environmental variables (Supplementary material, Table S4).

**Figure 4.**
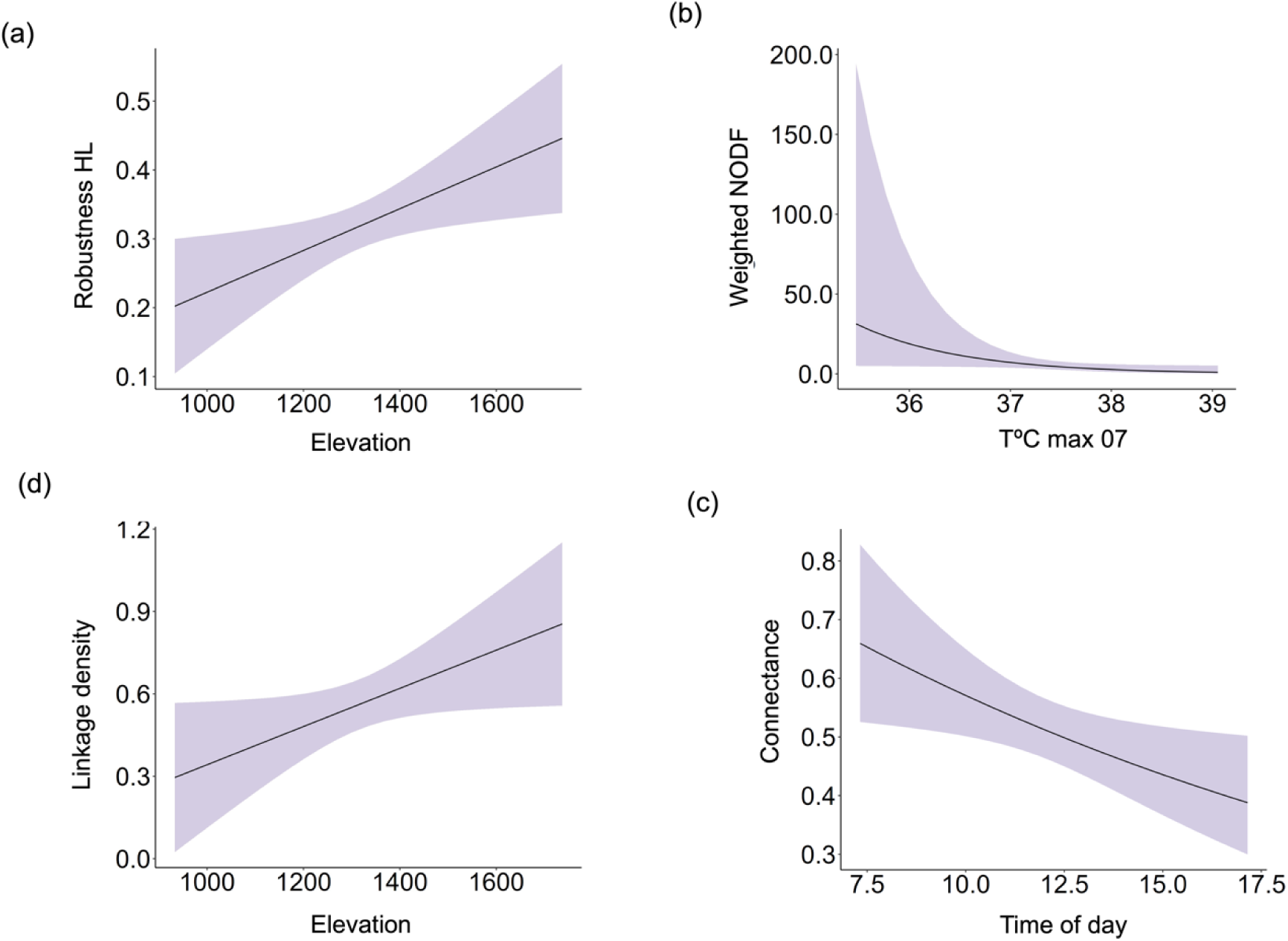
Modelled relationships based on generalized additive mixed models (GAMMs) of network descriptors vs elevation, microclimatic temperatures (T°C max), and the time of day. The tendencies are shown based on four samples each at 36 transects in 9 sites in the Sierra de Guadarrama.

**Table 1.** Summary of significance in Generalized Additive Mixed Models (GAMMs) testing the relationship between network descriptors and environmental variables, for 144 samples of trophic networks. Only significant response variables are shown for each model fitted with smoothing functions: elevation, date, time of day and maximum microclimatic temperature for June and July (Tmax_06 and Tmax_07). The random effects (re) include Site (9) and Transect (4 nested transects for each site).

| Response variable | Predictor | Edf<br>(smooth) | P-value |  | Adjusted<br>r<sup>2</sup> | Deviance<br>explained | Family | Link<br>function |
| --- | --- | --- | --- | --- | --- | --- | --- | --- |
| Robustness HL | Intercept | - | <0.001 | *** | 0.347 | 47 % | Gaussian | Identity |
|  | Elevation | 1.000 | 0.0166 | * |  |  |  |  |
|  | Transect [re] | 14.941 | 0.0037 | ** |  |  |  |  |
|  | Site [re] | <0.001 | 0.42447 | n. s |  |  |  |  |
| Weighted NODF | Intercept | - | <0.001 | *** | 0.476 | 46.2% | quasi-Poisson | log |
|  | Tmax_07 | 1.000 | 0.045 | * |  |  |  |  |
|  | Transect [re] | 10.43 | 0.118 | n. s |  |  |  |  |
|  | Site [re] | <0.001 | 0.645 | n. s |  |  |  |  |
| Connectance (Log) | Intercept | - | <0.001 | *** | 0.194 | 31.8% | Gaussian | Identity |
|  | Time of day | 1.000 | 0.0209 | * |  |  |  |  |
|  | Transect [re] | 12.427 | 0.0278 | * |  |  |  |  |
|  | Site [re] | <0.001 | 0.5987 | n. s |  |  |  |  |
| Linkage density | Intercept | - | <0.001 | *** | 0.202 | 31.5% | quasi-Poisson | log |
|  | Elevation | 1.000 | 0.0448 | * |  |  |  |  |
|  | Transect [re] | 6.797 | 0.1627 | n. s |  |  |  |  |
|  | Site [re] | 2.751 | 0.0698 | . |  |  |  |  |

In addition, for butterfly richness, the significant model (*n* = 144, r^2^adj = 0.543, Deviance explained = 54.3%, *p* = <0.001) revealed that butterfly richness at the sites during July increased linearly with elevation (edf = 1.000, *p* = 0.0048).

## DISCUSSION

We examined effects of elevation and microclimate on summer butterfly-flower interaction networks in a Mediterranean mountain system. At all sites, interactions were dominated by a few abundant butterfly and plant species, and network properties associated with stability were low at most sites. However, networks showed greater nestedness in sites with the coolest summer temperatures, and butterfly networks were more robust to the loss of flowering plant species at higher elevations. Our results suggest that habitat or microclimatic variation at high-elevation sites can increase the resilience of insect communities to climate change through their effects on properties of nestedness or modularity in insect-flower networks. These high-mountain ecosystems represent a priority for plant-pollinator conservation as climatic warming increasingly affects insect and plant communities in lower-elevation landscapes.

### Network variation over an elevation gradient

Networks in our system were dominated by interactions between a few butterfly and plant species (Fig. 1), although the identity of dominant species varied with elevation (see also Sánchez-Dávila et al., 2024). The proportion of possible links per site between insects and plants (connectance) did not vary with elevation, and we found no changes in the effective number of producers and consumers (vulnerability and generality). However, our analysis revealed that butterfly networks were more robust to the loss of flowering plant species at high elevations, related to an increase with elevation in the linkage density (the number of flower species visited per butterfly; and the number of butterfly species visiting each flower).

Network resilience to secondary extinctions is often linked to network compartmentalization and modularity (Memmott et al., 2004; Dalsgaard et al., 2013), and higher levels of nestedness can enhance resilience to disturbances, reduce interspecific competition and foster coexistence (Bastolla et al., 2009). We found greater nestedness in sites with the coldest modelled July temperatures, especially at the highest elevation where butterfly-flower interaction networks were more nested than elsewhere (1730 m; top left site panel in Fig. 1). At this site, butterfly interactions were concentrated among relatively few plant species, but modularity was provided because half of the butterfly species interacted with plants growing in damp pasture alongside a north-facing river valley (e.g., *Jacobaea vulgaris*), whilst the remaining half visited flower species growing on rockier south-facing terrain (e.g. *Carduus acanthoides*) (Figure 1).

### Factors influencing network properties

One specific way in which microclimates may influence observed network properties is through their effects on daily insect activity. In the hot and dry Mediterranean summer, nectar availability is a limiting factor for butterfly activity, providing both water and sugars (Lebeau et al., 2016a, b). Nectar depletion during the day likely influenced butterfly interactions, because as the day progressed network connectance decreased, meaning that butterflies interacted with a wider variety of flowers earlier in the morning. This pattern could have resulted from butterflies preferentially visiting plants with more dilute nectar later in the day (to avoid dehydration), or because they avoided visiting excessively hot flowers (Descamps et al., 2021) by limiting their foraging to plants growing in cooler microhabitats such as in the shade (see a counter example for advantages of foraging in warm flowers under cool conditions; Rands & Whitney, 2008). Either way, the result suggests diel changes to network topology due to heightened foraging costs during the hottest conditions in July. Temperatures in high elevation sites were cooler, leading either to longer nectar availability during the day, or to longer diurnal periods in which butterflies were able to nectar in open habitats and to forage from a larger number of available flowering plant species.

Flower-visitor networks vary through the season due to the phenology of component insects and plants (CaraDonna et al., 2017), responding to phenological delays at increased elevation for many butterflies (Gutiérrez Illán et al., 2012) and flowers (Bucher & Römermann, 2020). We found no effects of date on network properties, but this may be partly because June was cool and cloudy before the survey period (Fig S1), so that many flight and flowering periods were delayed before beginning together in July. At the high elevation sites, which warm up and dry out more slowly, the length of time over which flowers were maintained would have been longer. In these sites, plant species whose flowers were abundant and attractive to insects such as *R. ulmifolius*, *C. acanthoides*, or *J. vulgaris*, may have maintained nectar production with high sugar concentrations for longer (Gyan & Woodell, 1987; Giurfa & Núñez, 1992; Hicks et al., 2016), increasing network robustness relative to hotter, low elevation sites.

Dominant butterflies and plants varied across elevations, related to species’ elevation limits and habitat requirements (Gutiérrez et al., 2016). Butterfly species richness in the Sierra de Guadarrama peaks at mid-elevations, forming a humped elevation gradient (Gutiérrez Illán et al., 2010; Wilson et al., 2007). This happens because (i) the distribution and richness of butterfly species depends on the elevation ranges of their larval host plants (Gutiérrez et al., 2016), and (ii) the environmental variability and predominant forested vegetation at mid-elevations provides heterogeneous habitats allowing coexistence of generalist and specialist butterfly species (Álvarez et al., 2024). Over the past two decades, butterfly diversity in the Sierra de Guadarrama has declined in sites where early summer microclimates have warmed the most (Álvarez et al., 2024). Whilst these declines are probably related in part to effects of warming on butterfly juvenile stages (e.g., Merrill et al., 2006), our study indicates that possible impacts of climate change on nectar plant availability and adult foraging activity should also not be overlooked (see also WallisDeVries et al., 2012).

### Conservation implications

Our results suggest a role of local habitat heterogeneity in increasing plant-pollinator network resilience at cool, high elevations. The greater nestedness of the butterfly-flower network at the 1730 m site in July (Figure 1) is consistent with findings from the Cape Floristic Region, where generality and linkage density increased at intermediate elevations due to the presence of an ecotone (Adedoja et al., 2018). Ecotones, by providing local variation in habitat and floral species composition, can foster network stability and resilience by providing modularity. Maintaining local variation in habitat structure could therefore benefit insect communities by increasing the diversity of both larval and adult resources, including by increasing the robustness of nectar plant networks for flower-visiting species.

Our data were collected in one season of one year. Whilst we recorded interactions for more than half the butterfly species documented in the region (54 of 106 species, Álvarez et al., 2024), it is inevitable that network properties during different years or different seasons would vary (CaraDonna et al., 2017). Multiannual studies suggest that many plant-pollinator interactions are opportunistic (Petanidou et al., 2008), highlighting the need for studies across several years and seasons to assess how annual climate variability and the associated shifts in plant and insect phenology and abundance affect network dynamics. Nevertheless, our results are relevant for understanding factors influencing plant-pollinator networks in Mediterranean environments at a time of year when conditions of summer heat and drought can impose extreme limitations on activity and survival for flower-visiting insects. Our work suggests that understanding how habitat management or microclimate influence nectar availability or insect foraging costs under such conditions will become increasingly important to adapt insect conservation to a warming climate.

## Author contributions

Mario Alamo: Methodology; formal analysis; writing – original draft; writing – review and editing. Hugo Alejandro Álvarez: Conceptualization; funding acquisition; methodology; formal analysis; validation; writing – review and editing. Mario Mingarro: Data curation; software. Robert J. Wilson: Conceptualization; funding acquisition; methodology; writing – review and editing.

## Acknowledgments

We thank the official Master’s program "Biodiversity in Tropical Areas and its Conservation" and its sponsoring institutions: the Universidad Internacional Menéndez Pelayo and the Consejo Superior de Investigaciones Científicas (CSIC). Special thanks to Mario García París, Diego Gil Tapetado, and Anabel Perdices for their valuable comments on the manuscript. Thanks to Guim Ursul for assisting with field data collection and to Joseph Williamson for providing code for 3D plotting. Research permits were provided by the autonomous community of Madrid.

## DATA AVAILABILITY STATEMENT

The data that support the findings of this study are available on Figshare DOI: 10.6084/m9.figshare.27679122.

