## Supplementary material for "Effects of elevation and microclimatic temperatures on butterfly-flower interaction networks in a Mediterranean mountain range": Suplment_Material

**Electronic Supplementary Material: Additional details of methods and results**

Mario Alamo, Hugo A. Álvarez*, Mario Mingarro, Robert J. Wilson*

**Table S1** Summary of the 9 sites data. Plant Richness (PR), Butterfly Richness (BR), Nº Interactions (NI), mean maximum microclimatic temperature of the 4 transects per site (*n*=144 for June and July (mean Tmax 06 and mean Tmax 07), Top Interacting Butterfly Species (TIBS), Top Interacting Plant Species (TIPS) and Vegetation Series (VS). For habitat: Woodland edge (WE), Woodland clearing (WC), Woodland riverside (WR), Open woodland (OW) Scrubland (S), Grassland (G). Vegetation series based on Rivas-Martínez (1987) classification: 13a- Guadarrama Oromediterranean Silicicolous Series; 18a- Supramediterranean Carpetano-Iberian-Alcarreño subhumid Series and 24a- Supra-mesomediterranean Guadarrama, Iberian-Sorian, Celtiberian-Alcarreño, and Silicicolous Leonese Series. Plant and butterfly interactions are indicated in Brackets.

| Sites | PR | BR | Habitat | VS | NI | Mean Tmax 06 | Mean Tmax_07 | TIBS | TIPS |
|---|---|---|---|---|---|---|---|---|---|
| S1<br>1730 m | 7 | 11 | WE<br>(*Pinus sylvestris*) | 13a | 292 | 33.95 | 38.17 | *L. virgaureae* (116)<br>*H. lycaon*<br>(59) | *J. vulgaris*<br>(109)<br>*C. acanthoides* (105) |
| S2<br>1436 m | 11 | 11 | WE<br>WC<br>(*P. sylvestris*) | 18a | 73 | 34.27 | 38.72 | *M. lachesis* (18) | *C. acanthoides*<br>(30) |
| S3<br>1421 m | 15 | 21 | S<br>(*R. ulmifolius*)<br>WR<br>OW<br>(*P. sylvestris*) | 18a | 209 | 33.63 | 36.57 | *M. lachesis* (53)<br>*A. paphia*<br>(33) | *R. ulmifolius*<br>(120) |
| S4<br>1341 m | 11 | 14 | S<br>(*R. ulmifolius*)<br>*WC* | 18a | 153 | 36.21 | 38.85 | *A. pandora* (40)<br>*I. lathonia* (42) | *R. ulmifolius*<br>(77) |
| S5<br>1312 m | 15 | 13 | OW<br>WC | 18a | 77 | 34.11 | 38.21 | *M. lachesis* (16) | *E. campestre* (23)<br>*H. gouanii* (12) |
| S6<br>1250 m | 9 | 21 | OW<br>WC | 18a | 150 | 34.88 | 37.58 | *A. paphia* (35) | *R. ulmifolius* (101) |
| S7<br>1227 m | 12 | 12 | OW<br>(*Quercus pyrenaica*) | 18a | 68 | 33.45 | 37.65 | *M. lachesis* (18) | *J. montana* (11)<br>*C. alba* (12) |
| S8<br>1166 m | 18 | 13 | G<br>S<br>(*Rosa sp.*) | 18a | 55 | 34.49 | 36.87 | *P. tithonus* (15) | *E. campestre* (8) |
| S9<br>941 m | 14 | 12 | RS<br>OP<br>(Fraxinus sp.) | 24a | 68 | 34.45 | 35.59 | *P. tithonus* (20) | *E. campestre* (25)<br>*P. arabica* (17) |

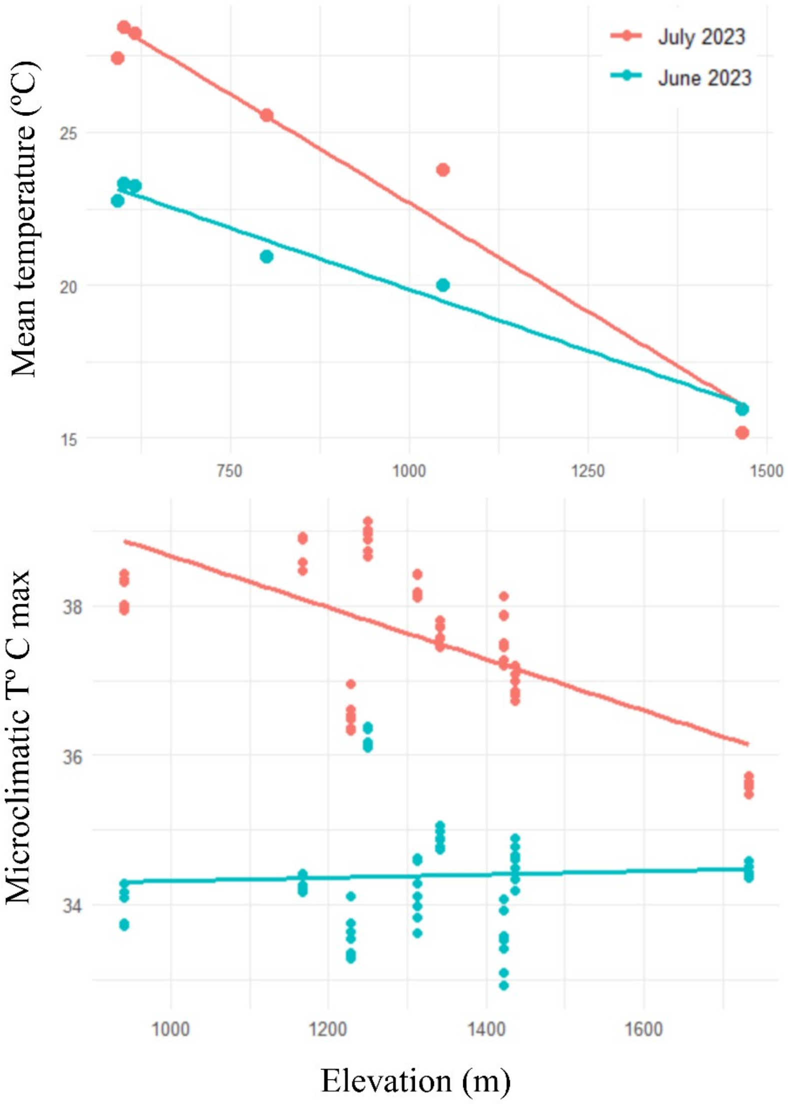


**Fig. S1** Scatterplot of: mean monthly temperature from 6 meteorological stations at different elevations in areas near the Sierra de Guadarrama (Station IDs: 8219099999 (Comenar Viejo), 8221099999 (Barajas), LEMD (Coslada), 8215099999 (Navacerrada), LETO (Torrejon de Ardoz), D4212 (La Cabrera)) (Top). Microclimatic maximum temperatures for June and July with elevation for the 36 transects (Down).

**Table S2** Summary of t-tests for Observed and mean Null Model (R2table method) values for 6 network descriptors: weighted NODF, generality, vulnerability, modularity Q, robustness HL, and robustness LL. Comparisons were conducted for a total of 144 matrices. Only those matrices with a sufficient number of interactions for statistical tests are shown. The first letters indicate the sampling point, the first number represents the transect, and the second number represents the visit. Observed: Observed value in the network. Null Mean: Mean null model value for 1000 random matrices per network. Lower CI: lower 95% confidence interval. Upper CI: upper 95% confidence interval. P-value: p-value of t-statistic. The first letters denote sites (S1-AM, S2-VI, S3-AA, S4-VII, S5-FR, S6-RLI, S7-HSA, S8-LA, S9-CR), where the first number represents the transect and the second number represents the visit.

| Network | Estimated Value | Weighted NODF | Generality | Vulnerability | Modularity Q | Robustness HL | Robustness LL |
|---|---|---|---|---|---|---|---|
| AA1.1 | Observed | 25 | 1.445 | 4.105 | 0.333 | 0.442 | 0.135 |
| | Null Mean | 24.695 | 2.454 | 2.454 | 0.245 | 0.405 | 0.405 |
| | Lower CI | 0 | 1.285 | 1.285 | 0.167 | 0.14 | 0.14 |
| | Upper CI | 33.333 | 4.117 | 4.117 | 0.389 | 0.698 | 0.698 |
| | t-Statistic | 91.86 | 109.551 | 109.551 | 123.13 | 72.04 | 72.04 |
| | P-Value | <0.001 | <0.001 | <0.001 | <0.001 | <0.001 | <0.001 |
| AA1.2 | Observed | 0 | 1.216 | 4.272 | 0.306 | 0.469 | 0.156 |
| | Null Mean | 23.971 | 2.889 | 2.889 | 0.203 | 0.413 | 0.413 |
| | Lower CI | 0 | 1.216 | 1.216 | 0.122 | 0.142 | 0.142 |
| | Upper CI | 41.667 | 4.828 | 4.828 | 0.306 | 0.707 | 0.707 |
| | t-Statistic | 51.867 | 86.129 | 86.129 | 118.831 | 74.795 | 74.795 |
| | P-Value | <0.001 | <0.001 | <0.001 | <0.001 | <0.001 | <0.001 |
| AA1.3 | Observed | 0 | 1 | 1.686 | 0.449 | 0.335 | 0.226 |
| | Null Mean | 15.544 | 2.024 | 2.024 | 0.33 | 0.335 | 0.335 |
| | Lower CI | 0 | 1.286 | 1.286 | 0.265 | 0.242 | 0.242 |
| | Upper CI | 33.333 | 2.992 | 2.992 | 0.449 | 0.416 | 0.416 |
| | t-Statistic | 33.956 | 144.886 | 144.886 | 200.767 | 267.271 | 267.271 |
| | P-Value | <0.001 | <0.001 | <0.001 | <0.001 | <0.001 | <0.001 |
| AA2.1 | Observed | 0 | 1 | 3.508 | 0.494 | 0.462 | 0.233 |
| | Null Mean | 0 | 2.561 | 2.561 | 0.428 | 0.397 | 0.397 |
| | Lower CI | 0 | 1 | 1 | 0.37 | 0.217 | 0.217 |
| | Upper CI | 0 | 4.556 | 4.556 | 0.494 | 0.569 | 0.569 |
| | t-Statistic | NA | 84.04 | 84.04 | 342.037 | 119.717 | 119.717 |
| | P-Value | NA | <0.001 | <0.001 | <0.001 | <0.001 | <0.001 |
| AA2.2 | Observed | 0 | 1 | 1.445 | 0.611 | 0.341 | 0.218 |
| | Null Mean | 0 | 1.777 | 1.777 | 0.427 | 0.357 | 0.357 |

| | | | | | | | |
|---|---|---|---|---|---|---|---|
| | Lower CI | 0 | 1 | 1 | 0.333 | 0.229 | 0.229 |
| | Upper CI | 0 | 2.333 | 2.333 | 0.611 | 0.456 | 0.456 |
| | t-Statistic | NA | 203.751 | 203.751 | 154.232 | 221.531 | 221.531 |
| | P-Value | NA | <0.001 | <0.001 | <0.001 | <0.001 | <0.001 |
| AA2.3 | Observed | 0 | 2.5 | 1 | 0.245 | 0.119 | 0.393 |
| | Null Mean | 17.871 | 2.152 | 2.152 | 0.199 | 0.285 | 0.285 |
| | Lower CI | 0 | 1 | 1 | 0.163 | 0.118 | 0.118 |
| | Upper CI | 42.857 | 3.383 | 3.383 | 0.245 | 0.458 | 0.458 |
| | t-Statistic | 26.731 | 101.598 | 101.598 | 185.331 | 91.38 | 91.38 |
| | P-Value | <0.001 | <0.001 | <0.001 | <0.001 | <0.001 | <0.001 |
| AA3.1 | Observed | 0 | 1 | 5.441 | 0.091 | 0.487 | 0.121 |
| | Null Mean | 15.655 | 3.558 | 3.558 | 0.077 | 0.311 | 0.311 |
| | Lower CI | 0 | 1 | 1 | 0.063 | 0.119 | 0.119 |
| | Upper CI | 27.586 | 6.274 | 6.274 | 0.091 | 0.526 | 0.526 |
| | t-Statistic | 42.402 | 65.049 | 65.049 | 236.267 | 79.895 | 79.895 |
| | P-Value | <0.001 | <0.001 | <0.001 | <0.001 | <0.001 | <0.001 |
| AA3.2 | Observed | 0 | 1 | 5.243 | 0.117 | 0.459 | 0.125 |
| | Null Mean | 13.59 | 3.456 | 3.456 | 0.1 | 0.309 | 0.309 |
| | Lower CI | 0 | 1 | 1 | 0.086 | 0.114 | 0.114 |
| | Upper CI | 24.138 | 6.184 | 6.184 | 0.117 | 0.524 | 0.524 |
| | t-Statistic | 35.875 | 65.683 | 65.683 | 242.842 | 79.485 | 79.485 |
| | P-Value | <0.001 | <0.001 | <0.001 | <0.001 | <0.001 | <0.001 |
| AA3.3 | Observed | 0 | 1 | 3.104 | 0.298 | 0.418 | 0.115 |
| | Null Mean | 17.564 | 2.61 | 2.61 | 0.206 | 0.372 | 0.372 |
| | Lower CI | 0 | 1 | 1 | 0.165 | 0.136 | 0.136 |
| | Upper CI | 31.818 | 4.158 | 4.158 | 0.298 | 0.625 | 0.625 |
| | t-Statistic | 36.595 | 91.548 | 91.548 | 157.59 | 79.097 | 79.097 |
| | P-Value | <0.001 | <0.001 | <0.001 | <0.001 | <0.001 | <0.001 |
| AA3.4 | Observed | 0 | 1 | 4.847 | 0.091 | 0.484 | 0.127 |
| | Null Mean | 17.932 | 3.248 | 3.248 | 0.075 | 0.31 | 0.31 |
| | Lower CI | 0 | 1 | 1 | 0.068 | 0.121 | 0.121 |
| | Upper CI | 27.273 | 5.541 | 5.541 | 0.091 | 0.517 | 0.517 |
| | t-Statistic | 47.311 | 68.287 | 68.287 | 268.843 | 81.688 | 81.688 |
| | P-Value | <0.001 | <0.001 | <0.001 | <0.001 | <0.001 | <0.001 |
| AA4.1 | Observed | 0 | 1 | 1.936 | 0.64 | 0.422 | 0.3 |
| | Null Mean | 4.503 | 2.254 | 2.254 | 0.44 | 0.429 | 0.429 |
| | Lower CI | 0 | 1.267 | 1.267 | 0.34 | 0.321 | 0.321 |
| | Upper CI | 14.286 | 3.6 | 3.6 | 0.58 | 0.523 | 0.523 |
| | t-Statistic | 28.297 | 143.564 | 143.564 | 221.368 | 270.512 | 270.512 |
| | P-Value | <0.001 | <0.001 | <0.001 | <0.001 | <0.001 | <0.001 |
| AA4.2 | Observed | 0 | 1 | 1.966 | 0.198 | 0.396 | 0.119 |
| | Null Mean | 37.543 | 1.856 | 1.856 | 0.117 | 0.283 | 0.283 |
| | Lower CI | 0 | 1 | 1 | 0.074 | 0.119 | 0.119 |

| | | | | | | | |
|---|---|---|---|---|---|---|---|
| | Upper CI | 57.143 | 2.712 | 2.712 | 0.198 | 0.457 | 0.457 |
| | t-Statistic | 43.744 | 124.6 | 124.6 | 62.88 | 90.702 | 90.702 |
| | P-Value | <0.001 | <0.001 | <0.001 | <0.001 | <0.001 | <0.001 |
| AA4.3 | Observed | 0 | 1.091 | 1.816 | 0.715 | 0.508 | 0.382 |
| | Null Mean | 13.25 | 2.942 | 2.942 | 0.431 | 0.488 | 0.488 |
| | Lower CI | 5.921 | 1.843 | 1.843 | 0.372 | 0.4 | 0.4 |
| | Upper CI | 19.408 | 4.351 | 4.351 | 0.512 | 0.58 | 0.58 |
| | t-Statistic | 112.316 | 197.259 | 197.259 | 368.005 | 335.473 | 335.473 |
| | P-Value | <0.001 | <0.001 | <0.001 | <0.001 | <0.001 | <0.001 |
| AA4.4 | Observed | 2.941 | 1.25 | 3.511 | 0.53 | 0.526 | 0.338 |
| | Null Mean | 18.577 | 3.498 | 3.498 | 0.277 | 0.527 | 0.527 |
| | Lower CI | 2.941 | 1.73 | 1.73 | 0.186 | 0.383 | 0.383 |
| | Upper CI | 32.846 | 5.702 | 5.702 | 0.385 | 0.648 | 0.648 |
| | t-Statistic | 74.754 | 102.956 | 102.956 | 163.848 | 236.203 | 236.203 |
| | P-Value | <0.001 | <0.001 | <0.001 | <0.001 | <0.001 | <0.001 |
| AM1.1 | Observed | 42.424 | 1.59 | 4.054 | 0.102 | 0.671 | 0.212 |
| | Null Mean | 30.121 | 2.626 | 2.626 | 0.177 | 0.402 | 0.402 |
| | Lower CI | 0 | 1.216 | 1.216 | 0.102 | 0.147 | 0.147 |
| | Upper CI | 63.636 | 4.094 | 4.094 | 0.276 | 0.68 | 0.68 |
| | t-Statistic | 52.547 | 99.978 | 99.978 | 99.991 | 79.431 | 79.431 |
| | P-Value | <0.001 | <0.001 | <0.001 | <0.001 | <0.001 | <0.001 |
| AM1.2 | Observed | 0 | 1 | 5.291 | 0.091 | 0.467 | 0.123 |
| | Null Mean | 10.043 | 3.463 | 3.463 | 0.071 | 0.311 | 0.311 |
| | Lower CI | 0 | 1 | 1 | 0.05 | 0.113 | 0.113 |
| | Upper CI | 21.739 | 6.217 | 6.217 | 0.091 | 0.531 | 0.531 |
| | t-Statistic | 29.29 | 65.208 | 65.208 | 114.082 | 77.767 | 77.767 |
| | P-Value | <0.001 | <0.001 | <0.001 | <0.001 | <0.001 | <0.001 |
| AM1.3 | Observed | 42.857 | 1.558 | 2.331 | 0.25 | 0.489 | 0.346 |
| | Null Mean | 52.128 | 2.3 | 2.3 | 0.112 | 0.453 | 0.453 |
| | Lower CI | 28.571 | 1.59 | 1.59 | 0.074 | 0.342 | 0.342 |
| | Upper CI | 71.429 | 3.074 | 3.074 | 0.18 | 0.572 | 0.572 |
| | t-Statistic | 137.26 | 188.512 | 188.512 | 124.405 | 275.489 | 275.489 |
| | P-Value | <0.001 | <0.001 | <0.001 | <0.001 | <0.001 | <0.001 |
| AM1.4 | Observed | 31.481 | 1.528 | 3.06 | 0.2 | 0.523 | 0.335 |
| | Null Mean | 52.903 | 2.607 | 2.607 | 0.127 | 0.468 | 0.468 |
| | Lower CI | 22.222 | 1.518 | 1.518 | 0.056 | 0.289 | 0.289 |
| | Upper CI | 72.222 | 3.82 | 3.82 | 0.202 | 0.653 | 0.653 |
| | t-Statistic | 130.941 | 124.196 | 124.196 | 107.134 | 149.113 | 149.113 |
| | P-Value | <0.001 | <0.001 | <0.001 | <0.001 | <0.001 | <0.001 |
| AM2.4 | Observed | 0 | 1 | 1.667 | 0.375 | 0.314 | 0.122 |
| | Null Mean | 0 | 1.658 | 1.658 | 0.314 | 0.248 | 0.248 |
| | Lower CI | 0 | 1 | 1 | 0.25 | 0.12 | 0.12 |
| | Upper CI | 0 | 2.5 | 2.5 | 0.375 | 0.403 | 0.403 |

| | | | | | | | |
|---|---|---|---|---|---|---|---|
| | t-Statistic | NA | 137.721 | 137.721 | 158.91 | 103.415 | 103.415 |
| | P-Value | NA | <0.001 | <0.001 | <0.001 | <0.001 | <0.001 |
| AM3.2 | Observed | 75 | 1.184 | 2.528 | 0.037 | 0.396 | 0.175 |
| | Null Mean | 60.9 | 1.822 | 1.822 | 0.05 | 0.278 | 0.278 |
| | Lower CI | 25 | 1.121 | 1.121 | 0.037 | 0.152 | 0.152 |
| | Upper CI | 75 | 2.528 | 2.528 | 0.079 | 0.408 | 0.408 |
| | t-Statistic | 106.374 | 124.129 | 124.129 | 94.927 | 111.445 | 111.445 |
| | P-Value | <0.001 | <0.001 | <0.001 | <0.001 | <0.001 | <0.001 |
| AM4.1 | Observed | 0 | 1 | 1.667 | 0.375 | 0.314 | 0.124 |
| | Null Mean | 0 | 1.66 | 1.66 | 0.311 | 0.249 | 0.249 |
| | Lower CI | 0 | 1 | 1 | 0.25 | 0.121 | 0.121 |
| | Upper CI | 0 | 2.5 | 2.5 | 0.375 | 0.402 | 0.402 |
| | t-Statistic | NA | 137.761 | 137.761 | 157.378 | 103.531 | 103.531 |
| | P-Value | NA | <0.001 | <0.001 | <0.001 | <0.001 | <0.001 |
| AM4.2 | Observed | 0 | 1 | 1.812 | 0.602 | 0.467 | 0.278 |
| | Null Mean | 16.562 | 2.599 | 2.599 | 0.359 | 0.455 | 0.455 |
| | Lower CI | 4.938 | 1.387 | 1.387 | 0.281 | 0.323 | 0.323 |
| | Upper CI | 25.926 | 4.09 | 4.09 | 0.469 | 0.588 | 0.588 |
| | t-Statistic | 94.135 | 140.799 | 140.799 | 243.42 | 202.279 | 202.279 |
| | P-Value | <0.001 | <0.001 | <0.001 | <0.001 | <0.001 | <0.001 |
| AM4.3 | Observed | 11.111 | 1.285 | 1.7 | 0.458 | 0.425 | 0.269 |
| | Null Mean | 22.807 | 2.335 | 2.335 | 0.254 | 0.447 | 0.447 |
| | Lower CI | 0 | 1.612 | 1.612 | 0.181 | 0.313 | 0.313 |
| | Upper CI | 38.889 | 3.008 | 3.008 | 0.389 | 0.569 | 0.569 |
| | t-Statistic | 55.33 | 248.716 | 248.716 | 141.738 | 264.586 | 264.586 |
| | P-Value | <0.001 | <0.001 | <0.001 | <0.001 | <0.001 | <0.001 |
| AM4.4 | Observed | 33.333 | 2.138 | 3.866 | 0.249 | 0.411 | 0.357 |
| | Null Mean | 41.646 | 2.574 | 2.574 | 0.192 | 0.444 | 0.444 |
| | Lower CI | 19.048 | 1.465 | 1.465 | 0.136 | 0.338 | 0.338 |
| | Upper CI | 61.905 | 3.875 | 3.875 | 0.286 | 0.552 | 0.552 |
| | t-Statistic | 98.09 | 139.168 | 139.168 | 146.309 | 277.721 | 277.721 |
| | P-Value | <0.001 | <0.001 | <0.001 | <0.001 | <0.001 | <0.001 |
| CR1.3 | Observed | 0 | 1.5 | 1.5 | 0.438 | 0.277 | 0.295 |
| | Null Mean | 0 | 1.425 | 1.425 | 0.477 | 0.273 | 0.273 |
| | Lower CI | 0 | 1 | 1 | 0.438 | 0.222 | 0.222 |
| | Upper CI | 0 | 1.5 | 1.5 | 0.625 | 0.299 | 0.299 |
| | t-Statistic | NA | 355.76 | 355.76 | 221.106 | 503.844 | 503.844 |
| | P-Value | NA | <0.001 | <0.001 | <0.001 | <0.001 | <0.001 |
| CR2.1 | Observed | 0 | 1.333 | 1.503 | 0.444 | 0.298 | 0.267 |
| | Null Mean | 15.983 | 1.825 | 1.825 | 0.314 | 0.316 | 0.316 |
| | Lower CI | 0 | 1.333 | 1.333 | 0.25 | 0.272 | 0.272 |
| | Upper CI | 33.333 | 2.219 | 2.219 | 0.444 | 0.364 | 0.364 |
| | t-Statistic | 34.074 | 256.623 | 256.623 | 151.818 | 481.722 | 481.722 |

| | | | | | | | |
|---|---|---|---|---|---|---|---|
| | P-Value | <0.001 | <0.001 | <0.001 | <0.001 | <0.001 | <0.001 |
| CR2.2 | Observed | 0 | 1.8 | 1 | 0.278 | 0.121 | 0.313 |
| | Null Mean | 24.85 | 1.847 | 1.847 | 0.204 | 0.268 | 0.268 |
| | Lower CI | 0 | 1 | 1 | 0.167 | 0.123 | 0.123 |
| | Upper CI | 50 | 2.56 | 2.56 | 0.278 | 0.408 | 0.408 |
| | t-Statistic | 31.418 | 149.089 | 149.089 | 154.574 | 107.69 | 107.69 |
| | P-Value | <0.001 | <0.001 | <0.001 | <0.001 | <0.001 | <0.001 |
| CR3.3 | Observed | 33.333 | 1.445 | 1.445 | 0.444 | 0.281 | 0.285 |
| | Null Mean | 5.4 | 1.842 | 1.842 | 0.34 | 0.341 | 0.341 |
| | Lower CI | 0 | 1.333 | 1.333 | 0.222 | 0.269 | 0.269 |
| | Upper CI | 33.333 | 2.333 | 2.333 | 0.5 | 0.409 | 0.409 |
| | t-Statistic | 17.746 | 273.698 | 273.698 | 122.662 | 391.118 | 391.118 |
| | P-Value | <0.001 | <0.001 | <0.001 | <0.001 | <0.001 | <0.001 |
| CR4.2 | Observed | 0 | 1 | 1.667 | 0.375 | 0.32 | 0.123 |
| | Null Mean | 0 | 1.653 | 1.653 | 0.317 | 0.25 | 0.25 |
| | Lower CI | 0 | 1 | 1 | 0.25 | 0.12 | 0.12 |
| | Upper CI | 0 | 2.5 | 2.5 | 0.375 | 0.404 | 0.404 |
| | t-Statistic | NA | 137.57 | 137.57 | 160.641 | 103.656 | 103.656 |
| | P-Value | NA | <0.001 | <0.001 | <0.001 | <0.001 | <0.001 |
| CR4.3 | Observed | 0 | 1 | 1.604 | 0.32 | 0.317 | 0.122 |
| | Null Mean | 44.85 | 1.716 | 1.716 | 0.224 | 0.254 | 0.254 |
| | Lower CI | 0 | 1 | 1 | 0.16 | 0.12 | 0.12 |
| | Upper CI | 75 | 2.463 | 2.463 | 0.32 | 0.405 | 0.405 |
| | t-Statistic | 38.55 | 144.421 | 144.421 | 90.351 | 104.416 | 104.416 |
| | P-Value | <0.001 | <0.001 | <0.001 | <0.001 | <0.001 | <0.001 |
| FR1.1 | Observed | 0 | 1.5 | 1.834 | 0.5 | 0.459 | 0.36 |
| | Null Mean | 0 | 1.833 | 1.833 | 0.472 | 0.422 | 0.422 |
| | Lower CI | 0 | 1.25 | 1.25 | 0.375 | 0.32 | 0.32 |
| | Upper CI | 0 | 2.25 | 2.25 | 0.625 | 0.496 | 0.496 |
| | t-Statistic | NA | 270.629 | 270.629 | 210.03 | 350.475 | 350.475 |
| | P-Value | NA | <0.001 | <0.001 | <0.001 | <0.001 | <0.001 |
| FR1.2 | Observed | 8.333 | 1.632 | 1.519 | 0.481 | 0.387 | 0.387 |
| | Null Mean | 6.65 | 2.13 | 2.13 | 0.379 | 0.418 | 0.418 |
| | Lower CI | 0 | 1.519 | 1.519 | 0.247 | 0.357 | 0.357 |
| | Upper CI | 25 | 3 | 3 | 0.543 | 0.472 | 0.472 |
| | t-Statistic | 27.987 | 288.686 | 288.686 | 163.824 | 615.38 | 615.38 |
| | P-Value | <0.001 | <0.001 | <0.001 | <0.001 | <0.001 | <0.001 |
| FR1.3 | Observed | 0 | 1.4 | 1.4 | 0.48 | 0.286 | 0.279 |
| | Null Mean | 0 | 1.643 | 1.643 | 0.387 | 0.321 | 0.321 |
| | Lower CI | 0 | 1 | 1 | 0.32 | 0.222 | 0.222 |
| | Upper CI | 0 | 1.8 | 1.8 | 0.64 | 0.359 | 0.359 |
| | t-Statistic | NA | 305.807 | 305.807 | 124.364 | 376.868 | 376.868 |
| | P-Value | NA | <0.001 | <0.001 | <0.001 | <0.001 | <0.001 |

| | | | | | | | |
|---|---|---|---|---|---|---|---|
| FR2.1 | Observed | 0 | 1.602 | 1.667 | 0.37 | 0.344 | 0.357 |
| | Null Mean | 39.242 | 1.973 | 1.973 | 0.224 | 0.356 | 0.356 |
| | Lower CI | 0 | 1.533 | 1.533 | 0.123 | 0.282 | 0.282 |
| | Upper CI | 75 | 2.389 | 2.389 | 0.37 | 0.414 | 0.414 |
| | t-Statistic | 45.931 | 340.016 | 340.016 | 100.17 | 463.684 | 463.684 |
| | P-Value | <0.001 | <0.001 | <0.001 | <0.001 | <0.001 | <0.001 |
| FR3.1 | Observed | 11.905 | 2.164 | 1.334 | 0.5 | 0.316 | 0.468 |
| | Null Mean | 5.398 | 1.945 | 1.945 | 0.51 | 0.391 | 0.391 |
| | Lower CI | 0 | 1 | 1 | 0.469 | 0.282 | 0.282 |
| | Upper CI | 14.286 | 2.75 | 2.75 | 0.656 | 0.49 | 0.49 |
| | t-Statistic | 26.22 | 160.093 | 160.093 | 345.081 | 232.903 | 232.903 |
| | P-Value | <0.001 | <0.001 | <0.001 | <0.001 | <0.001 | <0.001 |
| FR4.1 | Observed | 22.222 | 1.763 | 1.381 | 0.49 | 0.255 | 0.406 |
| | Null Mean | 5.856 | 1.957 | 1.957 | 0.38 | 0.374 | 0.374 |
| | Lower CI | 0 | 1.286 | 1.286 | 0.265 | 0.262 | 0.262 |
| | Upper CI | 22.222 | 2.714 | 2.714 | 0.612 | 0.491 | 0.491 |
| | t-Statistic | 25.423 | 221.313 | 221.313 | 139.871 | 229.086 | 229.086 |
| | P-Value | <0.001 | <0.001 | <0.001 | <0.001 | <0.001 | <0.001 |
| FR4.2 | Observed | 0 | 1.857 | 1.286 | 0.592 | 0.331 | 0.397 |
| | Null Mean | 0 | 1.725 | 1.725 | 0.52 | 0.385 | 0.385 |
| | Lower CI | 0 | 1.286 | 1.286 | 0.449 | 0.31 | 0.31 |
| | Upper CI | 0 | 2.143 | 2.143 | 0.654 | 0.451 | 0.451 |
| | t-Statistic | NA | 236.139 | 236.139 | 257.575 | 364.11 | 364.11 |
| | P-Value | NA | <0.001 | <0.001 | <0.001 | <0.001 | <0.001 |
| HSA1.1 | Observed | 33.333 | 1.445 | 2.219 | 0.333 | 0.393 | 0.258 |
| | Null Mean | 19.833 | 1.845 | 1.845 | 0.366 | 0.323 | 0.323 |
| | Lower CI | 0 | 1 | 1 | 0.333 | 0.219 | 0.219 |
| | Upper CI | 33.333 | 3 | 3 | 0.5 | 0.403 | 0.403 |
| | t-Statistic | 38.31 | 146.723 | 146.723 | 174.337 | 244.232 | 244.232 |
| | P-Value | <0.001 | <0.001 | <0.001 | <0.001 | <0.001 | <0.001 |
| HSA2.1 | Observed | 0 | 1.857 | 1.667 | 0.531 | 0.368 | 0.355 |
| | Null Mean | 2.783 | 1.843 | 1.843 | 0.448 | 0.38 | 0.38 |
| | Lower CI | 0 | 1.379 | 1.379 | 0.347 | 0.325 | 0.325 |
| | Upper CI | 16.667 | 2.143 | 2.143 | 0.612 | 0.419 | 0.419 |
| | t-Statistic | 16.938 | 306.765 | 306.765 | 191.969 | 700.271 | 700.271 |
| | P-Value | <0.001 | <0.001 | <0.001 | <0.001 | <0.001 | <0.001 |
| HSA2.2 | Observed | 12.5 | 1.725 | 1.829 | 0.42 | 0.343 | 0.463 |
| | Null Mean | 19.872 | 2.138 | 2.138 | 0.368 | 0.398 | 0.398 |
| | Lower CI | 0 | 1.4 | 1.4 | 0.3 | 0.32 | 0.32 |
| | Upper CI | 28.125 | 2.668 | 2.668 | 0.46 | 0.494 | 0.494 |
| | t-Statistic | 78.821 | 257.508 | 257.508 | 228.595 | 315.905 | 315.905 |
| | P-Value | <0.001 | <0.001 | <0.001 | <0.001 | <0.001 | <0.001 |
| HSA2.3 | Observed | 0 | 1.488 | 1.286 | 0.204 | 0.188 | 0.19 |

| | | | | | | | |
|---|---|---|---|---|---|---|---|
| | Null Mean | 71.7 | 1.548 | 1.548 | 0.114 | 0.187 | 0.187 |
| | Lower CI | 0 | 1.286 | 1.286 | 0.082 | 0.175 | 0.175 |
| | Upper CI | 100 | 1.763 | 1.763 | 0.204 | 0.199 | 0.199 |
| | t-Statistic | 50.309 | 403.896 | 403.896 | 66.742 | 1344.217 | 1344.217 |
| | P-Value | <0.001 | <0.001 | <0.001 | <0.001 | <0.001 | <0.001 |
| HSA3.1 | Observed | 1.754 | 1.828 | 2.478 | 0.568 | 0.523 | 0.456 |
| | Null Mean | 9.722 | 2.737 | 2.737 | 0.468 | 0.506 | 0.506 |
| | Lower CI | 1.754 | 1.988 | 1.988 | 0.393 | 0.44 | 0.44 |
| | Upper CI | 17.251 | 3.556 | 3.556 | 0.568 | 0.569 | 0.569 |
| | t-Statistic | 77.974 | 307.328 | 307.328 | 330.255 | 541.28 | 541.28 |
| | P-Value | <0.001 | <0.001 | <0.001 | <0.001 | <0.001 | <0.001 |
| HSA4.3 | Observed | 25 | 1.534 | 1.934 | 0.24 | 0.4 | 0.167 |
| | Null Mean | 14.55 | 1.719 | 1.719 | 0.289 | 0.279 | 0.279 |
| | Lower CI | 0 | 1 | 1 | 0.24 | 0.126 | 0.126 |
| | Upper CI | 25 | 2.2 | 2.2 | 0.48 | 0.422 | 0.422 |
| | t-Statistic | 37.295 | 247.47 | 247.47 | 123.667 | 106.237 | 106.237 |
| | P-Value | <0.001 | <0.001 | <0.001 | <0.001 | <0.001 | <0.001 |
| LA2.3 | Observed | 0 | 1.632 | 1.558 | 0.469 | 0.371 | 0.38 |
| | Null Mean | 10.679 | 2.109 | 2.109 | 0.358 | 0.413 | 0.413 |
| | Lower CI | 0 | 1.558 | 1.558 | 0.259 | 0.354 | 0.354 |
| | Upper CI | 25 | 2.778 | 2.778 | 0.519 | 0.467 | 0.467 |
| | t-Statistic | 40.046 | 289.534 | 289.534 | 151.954 | 619.362 | 619.362 |
| | P-Value | <0.001 | <0.001 | <0.001 | <0.001 | <0.001 | <0.001 |
| LA2.4 | Observed | 0 | 1.5 | 1.5 | 0.438 | 0.279 | 0.289 |
| | Null Mean | 0 | 1.413 | 1.413 | 0.48 | 0.274 | 0.274 |
| | Lower CI | 0 | 1 | 1 | 0.438 | 0.222 | 0.222 |
| | Upper CI | 0 | 1.5 | 1.5 | 0.625 | 0.3 | 0.3 |
| | t-Statistic | NA | 333.284 | 333.284 | 218.786 | 507.663 | 507.663 |
| | P-Value | NA | <0.001 | <0.001 | <0.001 | <0.001 | <0.001 |
| LA3.3 | Observed | 0 | 1.286 | 1.571 | 0.694 | 0.422 | 0.361 |
| | Null Mean | 0 | 1.407 | 1.407 | 0.696 | 0.383 | 0.383 |
| | Lower CI | 0 | 1 | 1 | 0.633 | 0.322 | 0.322 |
| | Upper CI | 0 | 1.571 | 1.571 | 0.776 | 0.431 | 0.431 |
| | t-Statistic | NA | 390.238 | 390.238 | 561.001 | 469.797 | 469.797 |
| | P-Value | NA | <0.001 | <0.001 | <0.001 | <0.001 | <0.001 |
| RLI2.1 | Observed | 0 | 1 | 2.263 | 0.133 | 0.394 | 0.135 |
| | Null Mean | 44.4 | 1.952 | 1.952 | 0.082 | 0.29 | 0.29 |
| | Lower CI | 0 | 1 | 1 | 0.061 | 0.124 | 0.124 |
| | Upper CI | 57.143 | 2.835 | 2.835 | 0.133 | 0.459 | 0.459 |
| | t-Statistic | 70.888 | 114.494 | 114.494 | 98.019 | 92.942 | 92.942 |
| | P-Value | <0.001 | <0.001 | <0.001 | <0.001 | <0.001 | <0.001 |
| RLI2.2 | Observed | 0 | 1 | 1.706 | 0.1 | 0.397 | 0.125 |
| | Null Mean | 45.771 | 1.6 | 1.6 | 0.038 | 0.291 | 0.291 |

| | | | | | | | |
|---|---|---|---|---|---|---|---|
| | Lower CI | 0 | 1 | 1 | 0.022 | 0.124 | 0.124 |
| | Upper CI | 57.143 | 2.08 | 2.08 | 0.1 | 0.463 | 0.463 |
| | t-Statistic | 63.412 | 166.904 | 166.904 | 39.375 | 93.379 | 93.379 |
| | P-Value | <0.001 | <0.001 | <0.001 | <0.001 | <0.001 | <0.001 |
| RLI2.3 | Observed | 0 | 1 | 1.89 | 0.5 | 0.471 | 0.125 |
| | Null Mean | 0 | 1.99 | 1.99 | 0.296 | 0.351 | 0.351 |
| | Lower CI | 0 | 1 | 1 | 0.167 | 0.125 | 0.125 |
| | Upper CI | 0 | 3 | 3 | 0.5 | 0.594 | 0.594 |
| | t-Statistic | NA | 131.207 | 131.207 | 74.494 | 81.945 | 81.945 |
| | P-Value | NA | <0.001 | <0.001 | <0.001 | <0.001 | <0.001 |
| RLI2.4 | Observed | 21.875 | 1.467 | 4.925 | 0.2 | 0.651 | 0.168 |
| | Null Mean | 9.456 | 2.787 | 2.787 | 0.25 | 0.376 | 0.376 |
| | Lower CI | 0 | 1 | 1 | 0.2 | 0.126 | 0.126 |
| | Upper CI | 21.875 | 4.925 | 4.925 | 0.32 | 0.645 | 0.645 |
| | t-Statistic | 30.132 | 80.331 | 80.331 | 196.417 | 73.754 | 73.754 |
| | P-Value | <0.001 | <0.001 | <0.001 | <0.001 | <0.001 | <0.001 |
| RLI3.1 | Observed | 0 | 1 | 3.007 | 0.43 | 0.526 | 0.227 |
| | Null Mean | 18.3 | 2.779 | 2.779 | 0.325 | 0.4 | 0.4 |
| | Lower CI | 0 | 1 | 1 | 0.248 | 0.224 | 0.224 |
| | Upper CI | 39.583 | 5.347 | 5.347 | 0.43 | 0.573 | 0.573 |
| | t-Statistic | 40.459 | 85.114 | 85.114 | 205.816 | 125.622 | 125.622 |
| | P-Value | <0.001 | <0.001 | <0.001 | <0.001 | <0.001 | <0.001 |
| RLI3.2 | Observed | 0 | 1 | 2.166 | 0.444 | 0.389 | 0.125 |
| | Null Mean | 15.381 | 2.278 | 2.278 | 0.219 | 0.378 | 0.378 |
| | Lower CI | 0 | 1.297 | 1.297 | 0.074 | 0.152 | 0.152 |
| | Upper CI | 38.095 | 3.52 | 3.52 | 0.444 | 0.625 | 0.625 |
| | t-Statistic | 31.864 | 130.897 | 130.897 | 70.418 | 86.928 | 86.928 |
| | P-Value | <0.001 | <0.001 | <0.001 | <0.001 | <0.001 | <0.001 |
| RLI3.3 | Observed | 0 | 1.305 | 1.667 | 0.451 | 0.471 | 0.254 |
| | Null Mean | 29.562 | 2.127 | 2.127 | 0.26 | 0.396 | 0.396 |
| | Lower CI | 7.692 | 1.503 | 1.503 | 0.222 | 0.259 | 0.259 |
| | Upper CI | 42.308 | 2.967 | 2.967 | 0.375 | 0.553 | 0.553 |
| | t-Statistic | 88.217 | 320.783 | 320.783 | 173.059 | 155.164 | 155.164 |
| | P-Value | <0.001 | <0.001 | <0.001 | <0.001 | <0.001 | <0.001 |
| RLI3.4 | Observed | 5.556 | 1.267 | 2.169 | 0.57 | 0.568 | 0.254 |
| | Null Mean | 6.131 | 2.254 | 2.254 | 0.412 | 0.453 | 0.453 |
| | Lower CI | 0 | 1.267 | 1.267 | 0.28 | 0.256 | 0.256 |
| | Upper CI | 16.667 | 3.4 | 3.4 | 0.57 | 0.655 | 0.655 |
| | t-Statistic | 34.517 | 163.886 | 163.886 | 173.4 | 127.024 | 127.024 |
| | P-Value | <0.001 | <0.001 | <0.001 | <0.001 | <0.001 | <0.001 |
| RLI4.1 | Observed | 11.905 | 1.246 | 5.293 | 0.249 | 0.481 | 0.329 |
| | Null Mean | 22.583 | 3.581 | 3.581 | 0.222 | 0.405 | 0.405 |
| | Lower CI | 0 | 1.141 | 1.141 | 0.183 | 0.282 | 0.282 |

| | | | | | | | |
|---|---|---|---|---|---|---|---|
| | Upper CI | 45.238 | 6.678 | 6.678 | 0.271 | 0.512 | 0.512 |
| | t-Statistic | 64.559 | 70.968 | 70.968 | 288.136 | 213.476 | 213.476 |
| | P-Value | <0.001 | <0.001 | <0.001 | <0.001 | <0.001 | <0.001 |
| RLI4.2 | Observed | 0 | 1 | 1.829 | 0.54 | 0.439 | 0.22 |
| | Null Mean | 26.129 | 2.288 | 2.288 | 0.314 | 0.404 | 0.404 |
| | Lower CI | 0 | 1.267 | 1.267 | 0.23 | 0.256 | 0.256 |
| | Upper CI | 46.154 | 3.524 | 3.524 | 0.47 | 0.545 | 0.545 |
| | t-Statistic | 62.026 | 151.523 | 151.523 | 154.361 | 170.254 | 170.254 |
| | P-Value | <0.001 | <0.001 | <0.001 | <0.001 | <0.001 | <0.001 |
| RLI4.4 | Observed | 0 | 1.4 | 1.902 | 0.64 | 0.442 | 0.374 |
| | Null Mean | 2.21 | 2.072 | 2.072 | 0.506 | 0.433 | 0.433 |
| | Lower CI | 0 | 1.4 | 1.4 | 0.42 | 0.369 | 0.369 |
| | Upper CI | 8 | 2.8 | 2.8 | 0.62 | 0.482 | 0.482 |
| | t-Statistic | 21.82 | 217.91 | 217.91 | 260.971 | 600.768 | 600.768 |
| | P-Value | <0.001 | <0.001 | <0.001 | <0.001 | <0.001 | <0.001 |
| VI1.1 | Observed | 0 | 1 | 1.667 | 0.375 | 0.314 | 0.129 |
| | Null Mean | 0 | 1.668 | 1.668 | 0.316 | 0.249 | 0.249 |
| | Lower CI | 0 | 1 | 1 | 0.25 | 0.12 | 0.12 |
| | Upper CI | 0 | 2.5 | 2.5 | 0.375 | 0.403 | 0.403 |
| | t-Statistic | NA | 138.016 | 138.016 | 159.815 | 103.537 | 103.537 |
| | P-Value | NA | <0.001 | <0.001 | <0.001 | <0.001 | <0.001 |
| VI1.3 | Observed | 11.111 | 1.638 | 3.007 | 0.248 | 0.459 | 0.351 |
| | Null Mean | 16.533 | 2.348 | 2.348 | 0.25 | 0.399 | 0.399 |
| | Lower CI | 0 | 1.364 | 1.364 | 0.19 | 0.291 | 0.291 |
| | Upper CI | 33.333 | 3.364 | 3.364 | 0.355 | 0.483 | 0.483 |
| | t-Statistic | 48.957 | 146.664 | 146.664 | 154.067 | 289.493 | 289.493 |
| | P-Value | <0.001 | <0.001 | <0.001 | <0.001 | <0.001 | <0.001 |
| VI1.4 | Observed | 0 | 1.333 | 3 | 0.417 | 0.411 | 0.28 |
| | Null Mean | 0 | 1.967 | 1.967 | 0.453 | 0.332 | 0.332 |
| | Lower CI | 0 | 1 | 1 | 0.417 | 0.211 | 0.211 |
| | Upper CI | 0 | 3 | 3 | 0.5 | 0.446 | 0.446 |
| | t-Statistic | NA | 109.067 | 109.067 | 357.107 | 165.325 | 165.325 |
| | P-Value | NA | <0.001 | <0.001 | <0.001 | <0.001 | <0.001 |
| VI2.1 | Observed | 16.667 | 1.267 | 3.797 | 0.38 | 0.551 | 0.234 |
| | Null Mean | 9.931 | 2.673 | 2.673 | 0.37 | 0.399 | 0.399 |
| | Lower CI | 0 | 1 | 1 | 0.3 | 0.219 | 0.219 |
| | Upper CI | 22.917 | 5.2 | 5.2 | 0.46 | 0.571 | 0.571 |
| | t-Statistic | 34.679 | 84.579 | 84.579 | 241.983 | 122.914 | 122.914 |
| | P-Value | <0.001 | <0.001 | <0.001 | <0.001 | <0.001 | <0.001 |
| VI2.2 | Observed | 0 | 1 | 1.679 | 0.497 | 0.428 | 0.315 |
| | Null Mean | 20.406 | 2.415 | 2.415 | 0.323 | 0.397 | 0.397 |
| | Lower CI | 12.903 | 1.273 | 1.273 | 0.284 | 0.331 | 0.331 |
| | Upper CI | 32.258 | 3.696 | 3.696 | 0.391 | 0.459 | 0.459 |

| | | | | | | | |
|---|---|---|---|---|---|---|---|
| | t-Statistic | 110.104 | 128.488 | 128.488 | 235.371 | 425.448 | 425.448 |
| | P-Value | <0.001 | <0.001 | <0.001 | <0.001 | <0.001 | <0.001 |
| VI2.3 | Observed | 0 | 1 | 1.503 | 0.5 | 0.338 | 0.216 |
| | Null Mean | 19.567 | 1.85 | 1.85 | 0.367 | 0.32 | 0.32 |
| | Lower CI | 0 | 1 | 1 | 0.333 | 0.218 | 0.218 |
| | Upper CI | 33.333 | 3 | 3 | 0.5 | 0.401 | 0.401 |
| | t-Statistic | 37.681 | 147.168 | 147.168 | 173.109 | 239.532 | 239.532 |
| | P-Value | <0.001 | <0.001 | <0.001 | <0.001 | <0.001 | <0.001 |
| VI2.4 | Observed | 0 | 1.8 | 1.4 | 0.48 | 0.267 | 0.408 |
| | Null Mean | 0 | 1.519 | 1.519 | 0.499 | 0.327 | 0.327 |
| | Lower CI | 0 | 1 | 1 | 0.44 | 0.219 | 0.219 |
| | Upper CI | 0 | 1.8 | 1.8 | 0.64 | 0.42 | 0.42 |
| | t-Statistic | NA | 264.55 | 264.55 | 204.521 | 206.313 | 206.313 |
| | P-Value | NA | <0.001 | <0.001 | <0.001 | <0.001 | <0.001 |
| VI3.1 | Observed | 0 | 1.667 | 1.667 | 0.222 | 0.188 | 0.173 |
| | Null Mean | 0 | 1.439 | 1.439 | 0.298 | 0.167 | 0.167 |
| | Lower CI | 0 | 1 | 1 | 0.222 | 0.125 | 0.125 |
| | Upper CI | 0 | 1.667 | 1.667 | 0.444 | 0.199 | 0.199 |
| | t-Statistic | NA | 203.629 | 203.629 | 89.405 | 249.181 | 249.181 |
| | P-Value | NA | <0.001 | <0.001 | <0.001 | <0.001 | <0.001 |
| VI4.2 | Observed | 0 | 1.667 | 1.667 | 0.222 | 0.188 | 0.186 |
| | Null Mean | 0 | 1.433 | 1.433 | 0.294 | 0.166 | 0.166 |
| | Lower CI | 0 | 1 | 1 | 0.222 | 0.125 | 0.125 |
| | Upper CI | 0 | 1.667 | 1.667 | 0.444 | 0.199 | 0.199 |
| | t-Statistic | NA | 201.311 | 201.311 | 89.414 | 246.966 | 246.966 |
| | P-Value | NA | <0.001 | <0.001 | <0.001 | <0.001 | <0.001 |
| VII1.1 | Observed | 0 | 1 | 1.563 | 0.521 | 0.409 | 0.229 |
| | Null Mean | 34.705 | 2.191 | 2.191 | 0.24 | 0.399 | 0.399 |
| | Lower CI | 15.385 | 1.464 | 1.464 | 0.172 | 0.256 | 0.256 |
| | Upper CI | 50 | 3.078 | 3.078 | 0.32 | 0.548 | 0.548 |
| | t-Statistic | 78.247 | 221.825 | 221.825 | 142.966 | 168.591 | 168.591 |
| | P-Value | <0.001 | <0.001 | <0.001 | <0.001 | <0.001 | <0.001 |
| VII1.2 | Observed | 0 | 1 | 3.298 | 0.508 | 0.544 | 0.211 |
| | Null Mean | 18.886 | 3.162 | 3.162 | 0.317 | 0.46 | 0.46 |
| | Lower CI | 3.226 | 1.314 | 1.314 | 0.238 | 0.257 | 0.257 |
| | Upper CI | 30.108 | 5.538 | 5.538 | 0.422 | 0.652 | 0.652 |
| | t-Statistic | 81.917 | 87.551 | 87.551 | 216.137 | 123.352 | 123.352 |
| | P-Value | <0.001 | <0.001 | <0.001 | <0.001 | <0.001 | <0.001 |
| VII1.3 | Observed | 0 | 1 | 2.118 | 0.165 | 0.401 | 0.13 |
| | Null Mean | 37.429 | 1.928 | 1.928 | 0.1 | 0.29 | 0.29 |
| | Lower CI | 0 | 1 | 1 | 0.066 | 0.124 | 0.124 |
| | Upper CI | 57.143 | 2.792 | 2.792 | 0.165 | 0.46 | 0.46 |
| | t-Statistic | 43.551 | 120.667 | 120.667 | 70.998 | 93.12 | 93.12 |

| | | | | | | | |
|---|---|---|---|---|---|---|---|
| | P-Value | <0.001 | <0.001 | <0.001 | <0.001 | <0.001 | <0.001 |
| VII1.4 | Observed | 0 | 1 | 2.534 | 0.18 | 0.414 | 0.13 |
| | Null Mean | 29.543 | 2.158 | 2.158 | 0.132 | 0.292 | 0.292 |
| | Lower CI | 0 | 1 | 1 | 0.1 | 0.126 | 0.126 |
| | Upper CI | 57.143 | 3.314 | 3.314 | 0.18 | 0.459 | 0.459 |
| | t-Statistic | 32.701 | 102.529 | 102.529 | 128.683 | 93.914 | 93.914 |
| | P-Value | <0.001 | <0.001 | <0.001 | <0.001 | <0.001 | <0.001 |
| VII2.1 | Observed | 0 | 1.445 | 1.445 | 0.5 | 0.291 | 0.285 |
| | Null Mean | 5.133 | 1.829 | 1.829 | 0.341 | 0.342 | 0.342 |
| | Lower CI | 0 | 1.333 | 1.333 | 0.222 | 0.271 | 0.271 |
| | Upper CI | 33.333 | 2.333 | 2.333 | 0.5 | 0.413 | 0.413 |
| | t-Statistic | 17.397 | 264.238 | 264.238 | 117.161 | 379.453 | 379.453 |
| | P-Value | <0.001 | <0.001 | <0.001 | <0.001 | <0.001 | <0.001 |
| VII2.2 | Observed | 0 | 1 | 4.914 | 0.18 | 0.424 | 0.118 |
| | Null Mean | 7.936 | 3.267 | 3.267 | 0.164 | 0.303 | 0.303 |
| | Lower CI | 0 | 1 | 1 | 0.14 | 0.113 | 0.113 |
| | Upper CI | 27.273 | 6.052 | 6.052 | 0.18 | 0.518 | 0.518 |
| | t-Statistic | 20.249 | 66.997 | 66.997 | 293.698 | 79.359 | 79.359 |
| | P-Value | <0.001 | <0.001 | <0.001 | <0.001 | <0.001 | <0.001 |
| VII2.3 | Observed | 25 | 1.49 | 2.667 | 0.277 | 0.399 | 0.434 |
| | Null Mean | 29.858 | 2.483 | 2.483 | 0.245 | 0.416 | 0.416 |
| | Lower CI | 5 | 1.49 | 1.49 | 0.201 | 0.372 | 0.372 |
| | Upper CI | 45 | 3.592 | 3.592 | 0.339 | 0.475 | 0.475 |
| | t-Statistic | 89.197 | 150.174 | 150.174 | 235.419 | 648.602 | 648.602 |
| | P-Value | <0.001 | <0.001 | <0.001 | <0.001 | <0.001 | <0.001 |
| VII3.1 | Observed | 16.667 | 1.573 | 1.342 | 0.34 | 0.333 | 0.328 |
| | Null Mean | 31.804 | 2.043 | 2.043 | 0.269 | 0.365 | 0.365 |
| | Lower CI | 0 | 1.342 | 1.342 | 0.22 | 0.323 | 0.323 |
| | Upper CI | 50 | 2.81 | 2.81 | 0.44 | 0.419 | 0.419 |
| | t-Statistic | 81.05 | 220.075 | 220.075 | 174.967 | 690.164 | 690.164 |
| | P-Value | <0.001 | <0.001 | <0.001 | <0.001 | <0.001 | <0.001 |
| VII3.2 | Observed | 0 | 1.333 | 1.828 | 0.604 | 0.436 | 0.395 |
| | Null Mean | 3.419 | 2.317 | 2.317 | 0.448 | 0.47 | 0.47 |
| | Lower CI | 0 | 1.612 | 1.612 | 0.347 | 0.401 | 0.401 |
| | Upper CI | 12 | 3.333 | 3.333 | 0.576 | 0.523 | 0.523 |
| | t-Statistic | 28.558 | 225.22 | 225.22 | 237.181 | 607.53 | 607.53 |
| | P-Value | <0.001 | <0.001 | <0.001 | <0.001 | <0.001 | <0.001 |
| VII3.3 | Observed | 0 | 1.627 | 1 | 0.625 | 0.224 | 0.444 |
| | Null Mean | 5.531 | 2.037 | 2.037 | 0.409 | 0.411 | 0.411 |
| | Lower CI | 0 | 1.25 | 1.25 | 0.313 | 0.255 | 0.255 |
| | Upper CI | 19.231 | 3 | 3 | 0.531 | 0.571 | 0.571 |
| | t-Statistic | 24.102 | 173.83 | 173.83 | 182.104 | 151.72 | 151.72 |
| | P-Value | <0.001 | <0.001 | <0.001 | <0.001 | <0.001 | <0.001 |

| | | | | | | | |
|---|---|---|---|---|---|---|---|
| VII4.1 | Observed | 0 | 1 | 1 | 0.406 | 0.222 | 0.222 |
| | Null Mean | 49.433 | 1.735 | 1.735 | 0.175 | 0.308 | 0.308 |
| | Lower CI | 0 | 1 | 1 | 0.125 | 0.222 | 0.222 |
| | Upper CI | 66.667 | 2.036 | 2.036 | 0.406 | 0.36 | 0.36 |
| | t-Statistic | 83.687 | 237.379 | 237.379 | 86.816 | 423.684 | 423.684 |
| | P-Value | <0.001 | <0.001 | <0.001 | <0.001 | <0.001 | <0.001 |
| VII4.2 | Observed | 19.231 | 1.232 | 1.764 | 0.521 | 0.504 | 0.27 |
| | Null Mean | 32.936 | 2.394 | 2.394 | 0.28 | 0.446 | 0.446 |
| | Lower CI | 11.538 | 1.643 | 1.643 | 0.189 | 0.291 | 0.291 |
| | Upper CI | 52.564 | 3.106 | 3.106 | 0.444 | 0.609 | 0.609 |
| | t-Statistic | 96.389 | 254.104 | 254.104 | 137.408 | 165.967 | 165.967 |

**Table S3** Summary of t-tests for Observed and mean Null Model (Vaznull method) values for 6 network descriptors: weighted NODF, generality, vulnerability, modularity Q, robustness HL, and robustness LL. Comparisons were conducted for a total of 144 matrices. Only those matrices with a sufficient number of interactions for statistical tests are shown. The first letters indicate the site, the first number represents the transect, and the second number represents the visit. Observed: Observed value in the network. Null Mean: Mean null model value for 1000 random matrices per network. Lower CI: lower 95% confidence interval. Upper CI: upper 95% confidence interval. P-value: p-value of t-statistic. The first letters denote the site (S1-AM, S2-VI, S3-AA, S4-VII, S5-FR, S6-RLI, S7-HSA, S8-LA, S9-CR), where the first number represents the transect and the second number represents the visit.

| Network | Estimated Value | weighted NODF | Generality | Vulnerability | Modularity Q | Robustness HL | Robustness LL |
|---|---|---|---|---|---|---|---|
| AA1.1 | Observed | 25.000 | 1.445 | 4.105 | 0.333 | 0.499 | 0.142 |
| | Null Mean | 18.460 | 2.442 | 2.442 | 0.281 | 0.379 | 0.379 |
| | Lower CI | 0.000 | 1.167 | 1.167 | 0.111 | 0.137 | 0.137 |
| | Upper CI | 37.500 | 4.709 | 4.709 | 0.417 | 0.669 | 0.669 |
| | t-Statistic | 46.892 | 91.312 | 91.312 | 108.121 | 71.312 | 71.312 |
| | P-Value | <0.001 | <0.001 | <0.001 | <0.001 | <0.001 | <0.001 |
| AA1.2 | Observed | 0.000 | 1.216 | 4.272 | 0.306 | 0.484 | 0.160 |
| | Null Mean | 16.786 | 2.650 | 2.650 | 0.217 | 0.360 | 0.360 |
| | Lower CI | 0.000 | 1.143 | 1.143 | 0.092 | 0.133 | 0.133 |
| | Upper CI | 37.500 | 5.215 | 5.215 | 0.408 | 0.665 | 0.665 |
| | t-Statistic | 44.337 | 79.914 | 79.914 | 77.143 | 72.743 | 72.743 |
| | P-Value | <0.001 | <0.001 | <0.001 | <0.001 | <0.001 | <0.001 |
| AA1.3 | Observed | 0.000 | 1.000 | 1.686 | 0.449 | 0.340 | 0.218 |
| | Null Mean | 12.239 | 1.901 | 1.901 | 0.379 | 0.340 | 0.340 |
| | Lower CI | 0.000 | 1.000 | 1.000 | 0.245 | 0.227 | 0.227 |
| | Upper CI | 55.556 | 2.992 | 2.992 | 0.571 | 0.451 | 0.451 |
| | t-Statistic | 24.213 | 153.919 | 153.919 | 137.509 | 240.087 | 240.087 |
| | P-Value | <0.001 | <0.001 | <0.001 | <0.001 | <0.001 | <0.001 |
| AA2.1 | Observed | 0.000 | 1.000 | 3.508 | 0.494 | 0.475 | 0.221 |
| | Null Mean | 1.714 | 2.471 | 2.471 | 0.439 | 0.393 | 0.393 |
| | Lower CI | 0.000 | 1.000 | 1.000 | 0.296 | 0.215 | 0.215 |
| | Upper CI | 25.000 | 4.713 | 4.713 | 0.593 | 0.609 | 0.609 |
| | t-Statistic | 9.624 | 80.576 | 80.576 | 154.534 | 117.376 | 117.376 |
| | P-Value | <0.001 | <0.001 | <0.001 | <0.001 | <0.001 | <0.001 |
| AA2.2 | Observed | 0.000 | 1.000 | 1.445 | 0.611 | 0.343 | 0.221 |
| | Null Mean | 2.922 | 1.749 | 1.749 | 0.444 | 0.336 | 0.336 |
| | Lower CI | 0.000 | 1.000 | 1.000 | 0.333 | 0.221 | 0.221 |
| | Upper CI | 33.333 | 3.000 | 3.000 | 0.611 | 0.454 | 0.454 |
| | t-Statistic | 11.507 | 169.403 | 169.403 | 170.393 | 216.639 | 216.639 |
| | P-Value | <0.001 | <0.001 | <0.001 | <0.001 | <0.001 | <0.001 |

| | | | | | | | |
|---|---|---|---|---|---|---|---|
| AA2.3 | Observed | 0.000 | 2.500 | 1.000 | 0.245 | 0.129 | 0.414 |
| | Null Mean | 14.843 | 2.050 | 2.050 | 0.241 | 0.289 | 0.289 |
| | Lower CI | 0.000 | 1.000 | 1.000 | 0.122 | 0.120 | 0.120 |
| | Upper CI | 57.143 | 3.383 | 3.383 | 0.490 | 0.527 | 0.527 |
| | t-Statistic | 22.153 | 102.312 | 102.312 | 87.118 | 87.951 | 87.951 |
| | P-Value | <0.001 | <0.001 | <0.001 | <0.001 | <0.001 | <0.001 |
| AA2.4 | Observed | 0.000 | 1.000 | 1.500 | 0.625 | 0.343 | 0.224 |
| | Null Mean | 0.000 | 1.731 | 1.731 | 0.462 | 0.332 | 0.332 |
| | Lower CI | 0.000 | 1.000 | 1.000 | 0.333 | 0.220 | 0.220 |
| | Upper CI | 0.000 | 3.000 | 3.000 | 0.625 | 0.451 | 0.451 |
| | t-Statistic | NA | 167.607 | 167.607 | 160.390 | 215.381 | 215.381 |
| | P-Value | NA | <0.001 | <0.001 | <0.001 | <0.001 | <0.001 |
| AA3.1 | Observed | 0.000 | 1.000 | 5.441 | 0.091 | 0.493 | 0.116 |
| | Null Mean | 13.537 | 3.617 | 3.617 | 0.097 | 0.317 | 0.317 |
| | Lower CI | 0.000 | 1.000 | 1.000 | 0.059 | 0.117 | 0.117 |
| | Upper CI | 27.586 | 7.016 | 7.016 | 0.204 | 0.638 | 0.638 |
| | t-Statistic | 39.228 | 63.412 | 63.412 | 77.879 | 76.875 | 76.875 |
| | P-Value | <0.001 | <0.001 | <0.001 | <0.001 | <0.001 | <0.001 |
| AA3.2 | Observed | 0.000 | 1.000 | 5.243 | 0.117 | 0.519 | 0.112 |
| | Null Mean | 10.613 | 3.607 | 3.607 | 0.130 | 0.322 | 0.322 |
| | Lower CI | 0.000 | 1.000 | 1.000 | 0.078 | 0.116 | 0.116 |
| | Upper CI | 27.586 | 6.930 | 6.930 | 0.305 | 0.646 | 0.646 |
| | t-Statistic | 30.096 | 63.548 | 63.548 | 77.177 | 75.337 | 75.337 |
| | P-Value | <0.001 | <0.001 | <0.001 | <0.001 | <0.001 | <0.001 |
| AA3.3 | Observed | 0.000 | 1.000 | 3.104 | 0.298 | 0.419 | 0.131 |
| | Null Mean | 11.618 | 2.278 | 2.278 | 0.231 | 0.314 | 0.314 |
| | Lower CI | 0.000 | 1.000 | 1.000 | 0.099 | 0.118 | 0.118 |
| | Upper CI | 45.455 | 4.405 | 4.405 | 0.463 | 0.594 | 0.594 |
| | t-Statistic | 22.855 | 83.587 | 83.587 | 68.027 | 77.547 | 77.547 |
| | P-Value | <0.001 | <0.001 | <0.001 | <0.001 | <0.001 | <0.001 |
| AA3.4 | Observed | 0.000 | 1.000 | 4.847 | 0.091 | 0.459 | 0.116 |
| | Null Mean | 16.142 | 3.241 | 3.241 | 0.092 | 0.314 | 0.314 |
| | Lower CI | 0.000 | 1.000 | 1.000 | 0.059 | 0.119 | 0.119 |
| | Upper CI | 31.818 | 6.195 | 6.195 | 0.190 | 0.616 | 0.616 |
| | t-Statistic | 41.194 | 67.013 | 67.013 | 79.923 | 78.617 | 78.617 |
| | P-Value | <0.001 | <0.001 | <0.001 | <0.001 | <0.001 | <0.001 |
| AA4.1 | Observed | 0.000 | 1.000 | 1.936 | 0.640 | 0.418 | 0.286 |
| | Null Mean | 5.748 | 1.989 | 1.989 | 0.508 | 0.403 | 0.403 |
| | Lower CI | 0.000 | 1.195 | 1.195 | 0.360 | 0.290 | 0.290 |
| | Upper CI | 23.810 | 3.597 | 3.597 | 0.660 | 0.520 | 0.520 |
| | t-Statistic | 24.733 | 132.093 | 132.093 | 196.938 | 236.845 | 236.845 |
| | P-Value | <0.001 | <0.001 | <0.001 | <0.001 | <0.001 | <0.001 |
| AA4.2 | Observed | 0.000 | 1.000 | 1.966 | 0.198 | 0.389 | 0.135 |
| | Null Mean | 31.664 | 2.058 | 2.058 | 0.174 | 0.293 | 0.293 |
| | Lower CI | 0.000 | 1.000 | 1.000 | 0.074 | 0.120 | 0.120 |
| | Upper CI | 57.143 | 3.442 | 3.442 | 0.346 | 0.528 | 0.528 |
| | t-Statistic | 40.533 | 109.720 | 109.720 | 68.563 | 89.007 | 89.007 |
| | P-Value | <0.001 | <0.001 | <0.001 | <0.001 | <0.001 | <0.001 |

| | | | | | | | |
|---|---|---|---|---|---|---|---|
| AA4.3 | Observed | 0.000 | 1.091 | 1.816 | 0.715 | 0.502 | 0.370 |
| | Null Mean | 8.381 | 2.544 | 2.544 | 0.549 | 0.476 | 0.476 |
| | Lower CI | 0.000 | 1.333 | 1.333 | 0.407 | 0.390 | 0.390 |
| | Upper CI | 18.429 | 4.568 | 4.568 | 0.711 | 0.576 | 0.576 |
| | t-Statistic | 55.342 | 127.456 | 127.456 | 212.546 | 334.846 | 334.846 |
| | P-Value | <0.001 | <0.001 | <0.001 | <0.001 | <0.001 | <0.001 |
| AA4.4 | Observed | 2.941 | 1.250 | 3.511 | 0.530 | 0.523 | 0.304 |
| | Null Mean | 13.631 | 2.441 | 2.441 | 0.430 | 0.440 | 0.440 |
| | Lower CI | 0.000 | 1.219 | 1.219 | 0.262 | 0.306 | 0.306 |
| | Upper CI | 28.934 | 4.570 | 4.570 | 0.561 | 0.595 | 0.595 |
| | t-Statistic | 60.007 | 96.906 | 96.906 | 172.441 | 172.289 | 172.289 |
| | P-Value | <0.001 | <0.001 | <0.001 | <0.001 | <0.001 | <0.001 |
| AM1.1 | Observed | 42.424 | 1.590 | 4.054 | 0.102 | 0.687 | 0.202 |
| | Null Mean | 29.294 | 2.785 | 2.785 | 0.162 | 0.437 | 0.437 |
| | Lower CI | 0.000 | 1.518 | 1.518 | 0.082 | 0.192 | 0.192 |
| | Upper CI | 63.636 | 4.332 | 4.332 | 0.265 | 0.686 | 0.686 |
| | t-Statistic | 57.350 | 107.620 | 107.620 | 106.818 | 82.412 | 82.412 |
| | P-Value | <0.001 | <0.001 | <0.001 | <0.001 | <0.001 | <0.001 |
| AM1.2 | Observed | 0.000 | 1.000 | 5.291 | 0.091 | 0.498 | 0.125 |
| | Null Mean | 10.984 | 4.185 | 4.185 | 0.099 | 0.326 | 0.326 |
| | Lower CI | 0.000 | 1.000 | 1.000 | 0.054 | 0.115 | 0.115 |
| | Upper CI | 21.739 | 8.481 | 8.481 | 0.204 | 0.667 | 0.667 |
| | t-Statistic | 34.920 | 59.799 | 59.799 | 74.721 | 73.602 | 73.602 |
| | P-Value | <0.001 | <0.001 | <0.001 | <0.001 | <0.001 | <0.001 |
| AM1.3 | Observed | 42.857 | 1.558 | 2.331 | 0.250 | 0.492 | 0.346 |
| | Null Mean | 39.795 | 2.327 | 2.327 | 0.154 | 0.415 | 0.415 |
| | Lower CI | 16.667 | 1.346 | 1.346 | 0.080 | 0.319 | 0.319 |
| | Upper CI | 54.762 | 3.853 | 3.853 | 0.284 | 0.532 | 0.532 |
| | t-Statistic | 121.181 | 133.751 | 133.751 | 85.654 | 286.595 | 286.595 |
| | P-Value | <0.001 | <0.001 | <0.001 | <0.001 | <0.001 | <0.001 |
| AM1.4 | Observed | 31.481 | 1.528 | 3.060 | 0.200 | 0.572 | 0.292 |
| | Null Mean | 44.918 | 2.650 | 2.650 | 0.157 | 0.436 | 0.436 |
| | Lower CI | 18.519 | 1.364 | 1.364 | 0.073 | 0.284 | 0.284 |
| | Upper CI | 64.815 | 4.486 | 4.486 | 0.269 | 0.613 | 0.613 |
| | t-Statistic | 113.137 | 105.950 | 105.950 | 99.054 | 153.968 | 153.968 |
| | P-Value | <0.001 | <0.001 | <0.001 | <0.001 | <0.001 | <0.001 |
| AM2.4 | Observed | 0.000 | 1.000 | 1.667 | 0.375 | 0.317 | 0.124 |
| | Null Mean | 0.000 | 1.713 | 1.713 | 0.310 | 0.257 | 0.257 |
| | Lower CI | 0.000 | 1.000 | 1.000 | 0.250 | 0.119 | 0.119 |
| | Upper CI | 0.000 | 2.500 | 2.500 | 0.500 | 0.415 | 0.415 |
| | t-Statistic | NA | 147.008 | 147.008 | 116.769 | 103.313 | 103.313 |
| | P-Value | NA | <0.001 | <0.001 | <0.001 | <0.001 | <0.001 |
| AM3.2 | Observed | 75.000 | 1.184 | 2.528 | 0.037 | 0.369 | 0.165 |
| | Null Mean | 57.475 | 1.840 | 1.840 | 0.067 | 0.278 | 0.278 |
| | Lower CI | 25.000 | 1.137 | 1.137 | 0.029 | 0.153 | 0.153 |
| | Upper CI | 75.000 | 2.781 | 2.781 | 0.140 | 0.410 | 0.410 |
| | t-Statistic | 87.758 | 126.799 | 126.799 | 68.577 | 111.507 | 111.507 |
| | P-Value | <0.001 | <0.001 | <0.001 | <0.001 | <0.001 | <0.001 |

| | | | | | | | |
|---|---|---|---|---|---|---|---|
| AM4.1 | Observed | 0.000 | 1.000 | 1.667 | 0.375 | 0.313 | 0.126 |
| | Null Mean | 0.000 | 1.706 | 1.706 | 0.312 | 0.256 | 0.256 |
| | Lower CI | 0.000 | 1.000 | 1.000 | 0.250 | 0.120 | 0.120 |
| | Upper CI | 0.000 | 2.500 | 2.500 | 0.500 | 0.412 | 0.412 |
| | t-Statistic | NA | 147.065 | 147.065 | 111.838 | 103.613 | 103.613 |
| | P-Value | NA | <0.001 | <0.001 | <0.001 | <0.001 | <0.001 |
| AM4.2 | Observed | 0.000 | 1.000 | 1.812 | 0.602 | 0.486 | 0.284 |
| | Null Mean | 10.707 | 2.234 | 2.234 | 0.452 | 0.418 | 0.418 |
| | Lower CI | 0.000 | 1.143 | 1.143 | 0.265 | 0.290 | 0.290 |
| | Upper CI | 29.630 | 4.272 | 4.272 | 0.643 | 0.564 | 0.564 |
| | t-Statistic | 36.526 | 109.239 | 109.239 | 143.066 | 192.913 | 192.913 |
| | P-Value | <0.001 | <0.001 | <0.001 | <0.001 | <0.001 | <0.001 |
| AM4.3 | Observed | 11.111 | 1.285 | 1.700 | 0.458 | 0.406 | 0.273 |
| | Null Mean | 19.622 | 1.808 | 1.808 | 0.350 | 0.344 | 0.344 |
| | Lower CI | 0.000 | 1.222 | 1.222 | 0.167 | 0.256 | 0.256 |
| | Upper CI | 55.556 | 2.930 | 2.930 | 0.514 | 0.458 | 0.458 |
| | t-Statistic | 37.803 | 177.816 | 177.816 | 104.664 | 231.401 | 231.401 |
| | P-Value | <0.001 | <0.001 | <0.001 | <0.001 | <0.001 | <0.001 |
| AM4.4 | Observed | 33.333 | 2.138 | 3.866 | 0.249 | 0.379 | 0.324 |
| | Null Mean | 32.907 | 2.555 | 2.555 | 0.238 | 0.421 | 0.421 |
| | Lower CI | 7.143 | 1.422 | 1.422 | 0.141 | 0.330 | 0.330 |
| | Upper CI | 54.762 | 4.446 | 4.446 | 0.401 | 0.532 | 0.532 |
| | t-Statistic | 87.396 | 115.844 | 115.844 | 111.758 | 292.866 | 292.866 |
| | P-Value | <0.001 | <0.001 | <0.001 | <0.001 | <0.001 | <0.001 |
| CR1.1 | Observed | 0.000 | 1.000 | 1.000 | 0.500 | 0.125 | 0.125 |
| | Null Mean | 0.000 | 1.443 | 1.443 | 0.315 | 0.166 | 0.166 |
| | Lower CI | 0.000 | 1.000 | 1.000 | 0.222 | 0.125 | 0.125 |
| | Upper CI | 0.000 | 1.667 | 1.667 | 0.500 | 0.199 | 0.199 |
| | t-Statistic | NA | 205.084 | 205.084 | 75.995 | 247.061 | 247.061 |
| | P-Value | NA | <0.001 | <0.001 | <0.001 | <0.001 | <0.001 |
| CR1.3 | Observed | 0.000 | 1.500 | 1.500 | 0.438 | 0.290 | 0.286 |
| | Null Mean | 0.000 | 1.683 | 1.683 | 0.404 | 0.305 | 0.305 |
| | Lower CI | 0.000 | 1.500 | 1.500 | 0.320 | 0.270 | 0.270 |
| | Upper CI | 0.000 | 2.200 | 2.200 | 0.500 | 0.359 | 0.359 |
| | t-Statistic | NA | 276.986 | 276.986 | 189.734 | 477.810 | 477.810 |
| | P-Value | NA | <0.001 | <0.001 | <0.001 | <0.001 | <0.001 |
| CR2.1 | Observed | 0.000 | 1.333 | 1.503 | 0.444 | 0.289 | 0.295 |
| | Null Mean | 14.550 | 1.737 | 1.737 | 0.350 | 0.310 | 0.310 |
| | Lower CI | 0.000 | 1.333 | 1.333 | 0.222 | 0.270 | 0.270 |
| | Upper CI | 66.667 | 2.219 | 2.219 | 0.500 | 0.361 | 0.361 |
| | t-Statistic | 24.476 | 244.559 | 244.559 | 129.734 | 472.203 | 472.203 |
| | P-Value | <0.001 | <0.001 | <0.001 | <0.001 | <0.001 | <0.001 |
| CR2.2 | Observed | 0.000 | 1.800 | 1.000 | 0.278 | 0.126 | 0.322 |
| | Null Mean | 22.200 | 1.720 | 1.720 | 0.238 | 0.262 | 0.262 |
| | Lower CI | 0.000 | 1.000 | 1.000 | 0.111 | 0.121 | 0.121 |
| | Upper CI | 75.000 | 2.560 | 2.560 | 0.444 | 0.411 | 0.411 |
| | t-Statistic | 25.396 | 147.460 | 147.460 | 82.270 | 105.499 | 105.499 |
| | P-Value | <0.001 | <0.001 | <0.001 | <0.001 | <0.001 | <0.001 |

| | | | | | | | |
|---|---|---|---|---|---|---|---|
| CR3.1 | Observed | 0.000 | 1.000 | 1.000 | 0.500 | 0.125 | 0.125 |
| | Null Mean | 0.000 | 1.448 | 1.448 | 0.324 | 0.166 | 0.166 |
| | Lower CI | 0.000 | 1.000 | 1.000 | 0.222 | 0.125 | 0.125 |
| | Upper CI | 0.000 | 1.667 | 1.667 | 0.500 | 0.199 | 0.199 |
| | t-Statistic | NA | 206.845 | 206.845 | 76.485 | 246.301 | 246.301 |
| | P-Value | NA | <0.001 | <0.001 | <0.001 | <0.001 | <0.001 |
| CR3.3 | Observed | 33.333 | 1.445 | 1.445 | 0.444 | 0.286 | 0.274 |
| | Null Mean | 11.917 | 1.663 | 1.663 | 0.378 | 0.306 | 0.306 |
| | Lower CI | 0.000 | 1.333 | 1.333 | 0.222 | 0.270 | 0.270 |
| | Upper CI | 66.667 | 2.219 | 2.219 | 0.500 | 0.359 | 0.359 |
| | t-Statistic | 22.227 | 252.532 | 252.532 | 140.098 | 472.083 | 472.083 |
| | P-Value | <0.001 | <0.001 | <0.001 | <0.001 | <0.001 | <0.001 |
| CR4.2 | Observed | 0.000 | 1.000 | 1.667 | 0.375 | 0.312 | 0.120 |
| | Null Mean | 0.000 | 1.713 | 1.713 | 0.306 | 0.256 | 0.256 |
| | Lower CI | 0.000 | 1.000 | 1.000 | 0.250 | 0.120 | 0.120 |
| | Upper CI | 0.000 | 2.500 | 2.500 | 0.500 | 0.413 | 0.413 |
| | t-Statistic | NA | 147.939 | 147.939 | 118.577 | 103.280 | 103.280 |
| | P-Value | NA | <0.001 | <0.001 | <0.001 | <0.001 | <0.001 |
| CR4.3 | Observed | 0.000 | 1.000 | 1.604 | 0.320 | 0.318 | 0.128 |
| | Null Mean | 19.975 | 1.727 | 1.727 | 0.269 | 0.261 | 0.261 |
| | Lower CI | 0.000 | 1.000 | 1.000 | 0.160 | 0.121 | 0.121 |
| | Upper CI | 75.000 | 2.463 | 2.463 | 0.480 | 0.413 | 0.413 |
| | t-Statistic | 19.850 | 152.970 | 152.970 | 90.029 | 104.796 | 104.796 |
| | P-Value | <0.001 | <0.001 | <0.001 | <0.001 | <0.001 | <0.001 |
| FR1.1 | Observed | 0.000 | 1.500 | 1.834 | 0.531 | 0.460 | 0.367 |
| | Null Mean | 3.497 | 1.904 | 1.904 | 0.479 | 0.400 | 0.400 |
| | Lower CI | 0.000 | 1.500 | 1.500 | 0.375 | 0.334 | 0.334 |
| | Upper CI | 21.875 | 2.743 | 2.743 | 0.547 | 0.474 | 0.474 |
| | t-Statistic | 17.207 | 244.654 | 244.654 | 334.858 | 376.126 | 376.126 |
| | P-Value | <0.001 | <0.001 | <0.001 | <0.001 | <0.001 | <0.001 |
| FR1.2 | Observed | 8.333 | 1.632 | 1.519 | 0.481 | 0.384 | 0.373 |
| | Null Mean | 11.804 | 1.814 | 1.814 | 0.436 | 0.375 | 0.375 |
| | Lower CI | 0.000 | 1.517 | 1.517 | 0.296 | 0.342 | 0.342 |
| | Upper CI | 37.500 | 2.550 | 2.550 | 0.556 | 0.423 | 0.423 |
| | t-Statistic | 32.401 | 279.992 | 279.992 | 183.694 | 847.041 | 847.041 |
| | P-Value | <0.001 | <0.001 | <0.001 | <0.001 | <0.001 | <0.001 |
| FR1.3 | Observed | 0.000 | 1.400 | 1.400 | 0.480 | 0.283 | 0.286 |
| | Null Mean | 4.800 | 1.644 | 1.644 | 0.395 | 0.303 | 0.303 |
| | Lower CI | 0.000 | 1.400 | 1.400 | 0.320 | 0.271 | 0.271 |
| | Upper CI | 33.333 | 2.200 | 2.200 | 0.480 | 0.359 | 0.359 |
| | t-Statistic | 12.964 | 263.625 | 263.625 | 171.653 | 477.145 | 477.145 |
| | P-Value | <0.001 | <0.001 | <0.001 | <0.001 | <0.001 | <0.001 |
| FR2.1 | Observed | 0.000 | 1.602 | 1.667 | 0.370 | 0.345 | 0.323 |
| | Null Mean | 29.883 | 1.907 | 1.907 | 0.256 | 0.340 | 0.340 |
| | Lower CI | 0.000 | 1.558 | 1.558 | 0.148 | 0.310 | 0.310 |
| | Upper CI | 66.667 | 2.510 | 2.510 | 0.395 | 0.368 | 0.368 |
| | t-Statistic | 41.004 | 327.368 | 327.368 | 111.146 | 1070.141 | 1070.141 |
| | P-Value | <0.001 | <0.001 | <0.001 | <0.001 | <0.001 | <0.001 |

| | | | | | | | |
|---|---|---|---|---|---|---|---|
| FR3.1 | Observed | 11.905 | 2.164 | 1.334 | 0.500 | 0.309 | 0.475 |
| | Null Mean | 1.026 | 2.024 | 2.024 | 0.515 | 0.404 | 0.404 |
| | Lower CI | 0.000 | 1.250 | 1.250 | 0.407 | 0.302 | 0.302 |
| | Upper CI | 14.286 | 3.500 | 3.500 | 0.625 | 0.523 | 0.523 |
| | t-Statistic | 8.911 | 135.830 | 135.830 | 253.524 | 235.261 | 235.261 |
| | P-Value | <0.001 | <0.001 | <0.001 | <0.001 | <0.001 | <0.001 |
| FR4.1 | Observed | 22.222 | 1.763 | 1.381 | 0.490 | 0.261 | 0.408 |
| | Null Mean | 9.350 | 1.782 | 1.782 | 0.410 | 0.345 | 0.345 |
| | Lower CI | 0.000 | 1.286 | 1.286 | 0.265 | 0.256 | 0.256 |
| | Upper CI | 38.889 | 2.992 | 2.992 | 0.531 | 0.456 | 0.456 |
| | t-Statistic | 23.444 | 191.463 | 191.463 | 172.925 | 227.996 | 227.996 |
| | P-Value | <0.001 | <0.001 | <0.001 | <0.001 | <0.001 | <0.001 |
| FR4.2 | Observed | 0.000 | 1.857 | 1.286 | 0.592 | 0.315 | 0.397 |
| | Null Mean | 0.850 | 1.831 | 1.831 | 0.512 | 0.385 | 0.385 |
| | Lower CI | 0.000 | 1.286 | 1.286 | 0.391 | 0.311 | 0.311 |
| | Upper CI | 12.500 | 2.750 | 2.750 | 0.653 | 0.469 | 0.469 |
| | t-Statistic | 8.160 | 192.927 | 192.927 | 236.946 | 346.495 | 346.495 |
| | P-Value | <0.001 | <0.001 | <0.001 | <0.001 | <0.001 | <0.001 |
| HSA1.1 | Observed | 33.333 | 1.445 | 2.219 | 0.333 | 0.367 | 0.266 |
| | Null Mean | 4.878 | 1.912 | 1.912 | 0.399 | 0.343 | 0.343 |
| | Lower CI | 0.000 | 1.333 | 1.333 | 0.333 | 0.256 | 0.256 |
| | Upper CI | 33.333 | 3.000 | 3.000 | 0.500 | 0.452 | 0.452 |
| | t-Statistic | 14.264 | 171.752 | 171.752 | 200.257 | 241.116 | 241.116 |
| | P-Value | <0.001 | <0.001 | <0.001 | <0.001 | <0.001 | <0.001 |
| HSA1.2 | Observed | 0.000 | 1.000 | 1.000 | 0.500 | 0.125 | 0.125 |
| | Null Mean | 0.000 | 1.432 | 1.432 | 0.310 | 0.167 | 0.167 |
| | Lower CI | 0.000 | 1.000 | 1.000 | 0.222 | 0.125 | 0.125 |
| | Upper CI | 0.000 | 1.667 | 1.667 | 0.500 | 0.199 | 0.199 |
| | t-Statistic | NA | 201.086 | 201.086 | 75.868 | 252.349 | 252.349 |
| | P-Value | NA | <0.001 | <0.001 | <0.001 | <0.001 | <0.001 |
| HSA2.1 | Observed | 0.000 | 1.857 | 1.667 | 0.531 | 0.394 | 0.360 |
| | Null Mean | 4.667 | 1.853 | 1.853 | 0.451 | 0.374 | 0.374 |
| | Lower CI | 0.000 | 1.571 | 1.571 | 0.367 | 0.341 | 0.341 |
| | Upper CI | 25.000 | 2.714 | 2.714 | 0.531 | 0.420 | 0.420 |
| | t-Statistic | 17.821 | 298.669 | 298.669 | 270.135 | 873.599 | 873.599 |
| | P-Value | <0.001 | <0.001 | <0.001 | <0.001 | <0.001 | <0.001 |
| HSA2.2 | Observed | 12.500 | 1.725 | 1.829 | 0.420 | 0.344 | 0.445 |
| | Null Mean | 14.156 | 2.056 | 2.056 | 0.419 | 0.398 | 0.398 |
| | Lower CI | 0.000 | 1.467 | 1.467 | 0.290 | 0.329 | 0.329 |
| | Upper CI | 37.500 | 3.000 | 3.000 | 0.560 | 0.487 | 0.487 |
| | t-Statistic | 39.057 | 227.565 | 227.565 | 181.140 | 356.439 | 356.439 |
| | P-Value | <0.001 | <0.001 | <0.001 | <0.001 | <0.001 | <0.001 |
| HSA2.3 | Observed | 0.000 | 1.488 | 1.286 | 0.204 | 0.179 | 0.191 |
| | Null Mean | 61.250 | 1.582 | 1.582 | 0.134 | 0.187 | 0.187 |
| | Lower CI | 0.000 | 1.286 | 1.286 | 0.041 | 0.175 | 0.175 |
| | Upper CI | 100.000 | 1.857 | 1.857 | 0.327 | 0.200 | 0.200 |
| | t-Statistic | 49.804 | 423.953 | 423.953 | 61.454 | 1327.069 | 1327.069 |
| | P-Value | <0.001 | <0.001 | <0.001 | <0.001 | <0.001 | <0.001 |

| | | | | | | | |
|---|---|---|---|---|---|---|---|
| HSA3.1 | Observed | 1.754 | 1.828 | 2.478 | 0.568 | 0.527 | 0.450 |
| | Null Mean | 8.408 | 2.380 | 2.380 | 0.544 | 0.488 | 0.488 |
| | Lower CI | 0.000 | 1.686 | 1.686 | 0.432 | 0.425 | 0.425 |
| | Upper CI | 18.432 | 3.493 | 3.493 | 0.632 | 0.557 | 0.557 |
| | t-Statistic | 53.248 | 223.408 | 223.408 | 326.262 | 517.527 | 517.527 |
| | P-Value | <0.001 | <0.001 | <0.001 | <0.001 | <0.001 | <0.001 |
| HSA3.2 | Observed | 0.000 | 1.667 | 1.000 | 0.444 | 0.123 | 0.313 |
| | Null Mean | 0.000 | 1.653 | 1.653 | 0.335 | 0.253 | 0.253 |
| | Lower CI | 0.000 | 1.000 | 1.000 | 0.250 | 0.119 | 0.119 |
| | Upper CI | 0.000 | 2.500 | 2.500 | 0.444 | 0.413 | 0.413 |
| | t-Statistic | NA | 149.525 | 149.525 | 109.704 | 102.563 | 102.563 |
| | P-Value | NA | <0.001 | <0.001 | <0.001 | <0.001 | <0.001 |
| HSA4.3 | Observed | 25.000 | 1.534 | 1.934 | 0.240 | 0.396 | 0.167 |
| | Null Mean | 20.475 | 1.828 | 1.828 | 0.252 | 0.282 | 0.282 |
| | Lower CI | 0.000 | 1.400 | 1.400 | 0.160 | 0.157 | 0.157 |
| | Upper CI | 75.000 | 2.463 | 2.463 | 0.320 | 0.420 | 0.420 |
| | t-Statistic | 23.707 | 209.322 | 209.322 | 142.009 | 108.629 | 108.629 |
| | P-Value | <0.001 | <0.001 | <0.001 | <0.001 | <0.001 | <0.001 |
| LA1.1 | Observed | 0.000 | 2.000 | 1.000 | 0.500 | 0.125 | 0.471 |
| | Null Mean | 0.000 | 1.838 | 1.838 | 0.376 | 0.299 | 0.299 |
| | Lower CI | 0.000 | 1.000 | 1.000 | 0.240 | 0.119 | 0.119 |
| | Upper CI | 0.000 | 3.400 | 3.400 | 0.500 | 0.537 | 0.537 |
| | t-Statistic | NA | 109.809 | 109.809 | 135.425 | 80.874 | 80.874 |
| | P-Value | NA | <0.001 | <0.001 | <0.001 | <0.001 | <0.001 |
| LA2.1 | Observed | 0.000 | 1.000 | 1.000 | 0.667 | 0.222 | 0.222 |
| | Null Mean | 0.000 | 1.605 | 1.605 | 0.437 | 0.296 | 0.296 |
| | Lower CI | 0.000 | 1.000 | 1.000 | 0.320 | 0.222 | 0.222 |
| | Upper CI | 0.000 | 2.200 | 2.200 | 0.667 | 0.357 | 0.357 |
| | t-Statistic | NA | 227.196 | 227.196 | 136.302 | 381.034 | 381.034 |
| | P-Value | NA | <0.001 | <0.001 | <0.001 | <0.001 | <0.001 |
| LA2.2 | Observed | 0.000 | 1.000 | 1.000 | 0.500 | 0.125 | 0.125 |
| | Null Mean | 0.000 | 1.433 | 1.433 | 0.312 | 0.165 | 0.165 |
| | Lower CI | 0.000 | 1.000 | 1.000 | 0.222 | 0.125 | 0.125 |
| | Upper CI | 0.000 | 1.667 | 1.667 | 0.500 | 0.199 | 0.199 |
| | t-Statistic | NA | 201.537 | 201.537 | 75.911 | 242.365 | 242.365 |
| | P-Value | NA | <0.001 | <0.001 | <0.001 | <0.001 | <0.001 |
| LA2.3 | Observed | 0.000 | 1.632 | 1.558 | 0.469 | 0.383 | 0.388 |
| | Null Mean | 11.925 | 1.849 | 1.849 | 0.424 | 0.375 | 0.375 |
| | Lower CI | 0.000 | 1.519 | 1.519 | 0.272 | 0.341 | 0.341 |
| | Upper CI | 37.500 | 2.643 | 2.643 | 0.556 | 0.422 | 0.422 |
| | t-Statistic | 31.150 | 268.335 | 268.335 | 174.817 | 815.018 | 815.018 |
| | P-Value | <0.001 | <0.001 | <0.001 | <0.001 | <0.001 | <0.001 |
| LA2.4 | Observed | 0.000 | 1.500 | 1.500 | 0.438 | 0.283 | 0.274 |
| | Null Mean | 0.000 | 1.688 | 1.688 | 0.405 | 0.305 | 0.305 |
| | Lower CI | 0.000 | 1.500 | 1.500 | 0.320 | 0.270 | 0.270 |
| | Upper CI | 0.000 | 2.200 | 2.200 | 0.500 | 0.362 | 0.362 |
| | t-Statistic | NA | 272.245 | 272.245 | 187.035 | 466.807 | 466.807 |
| | P-Value | NA | <0.001 | <0.001 | <0.001 | <0.001 | <0.001 |

| | | | | | | | |
|---|---|---|---|---|---|---|---|
| LA3.1 | Observed | 0.000 | 1.000 | 2.500 | 0.375 | 0.397 | 0.130 |
| | Null Mean | 0.000 | 1.978 | 1.978 | 0.344 | 0.292 | 0.292 |
| | Lower CI | 0.000 | 1.000 | 1.000 | 0.240 | 0.118 | 0.118 |
| | Upper CI | 0.000 | 3.400 | 3.400 | 0.500 | 0.531 | 0.531 |
| | t-Statistic | NA | 101.881 | 101.881 | 119.379 | 83.907 | 83.907 |
| | P-Value | NA | <0.001 | <0.001 | <0.001 | <0.001 | <0.001 |
| LA3.3 | Observed | 0.000 | 1.286 | 1.571 | 0.694 | 0.413 | 0.364 |
| | Null Mean | 0.000 | 1.822 | 1.822 | 0.592 | 0.414 | 0.414 |
| | Lower CI | 0.000 | 1.286 | 1.286 | 0.460 | 0.347 | 0.347 |
| | Upper CI | 0.000 | 2.556 | 2.556 | 0.735 | 0.489 | 0.489 |
| | t-Statistic | NA | 209.931 | 209.931 | 264.053 | 459.852 | 459.852 |
| | P-Value | NA | <0.001 | <0.001 | <0.001 | <0.001 | <0.001 |
| LA4.4 | Observed | 0.000 | 1.000 | 1.000 | 0.500 | 0.125 | 0.125 |
| | Null Mean | 0.000 | 1.438 | 1.438 | 0.311 | 0.167 | 0.167 |
| | Lower CI | 0.000 | 1.000 | 1.000 | 0.222 | 0.125 | 0.125 |
| | Upper CI | 0.000 | 1.667 | 1.667 | 0.500 | 0.199 | 0.199 |
| | t-Statistic | NA | 203.155 | 203.155 | 75.888 | 249.330 | 249.330 |
| | P-Value | NA | <0.001 | <0.001 | <0.001 | <0.001 | <0.001 |
| RLI2.1 | Observed | 0.000 | 1.000 | 2.263 | 0.133 | 0.386 | 0.119 |
| | Null Mean | 37.421 | 2.048 | 2.048 | 0.120 | 0.292 | 0.292 |
| | Lower CI | 0.000 | 1.000 | 1.000 | 0.051 | 0.120 | 0.120 |
| | Upper CI | 57.143 | 3.500 | 3.500 | 0.245 | 0.518 | 0.518 |
| | t-Statistic | 53.544 | 104.186 | 104.186 | 64.158 | 90.675 | 90.675 |
| | P-Value | <0.001 | <0.001 | <0.001 | <0.001 | <0.001 | <0.001 |
| RLI2.2 | Observed | 0.000 | 1.000 | 1.706 | 0.100 | 0.389 | 0.120 |
| | Null Mean | 45.957 | 1.836 | 1.836 | 0.069 | 0.294 | 0.294 |
| | Lower CI | 0.000 | 1.000 | 1.000 | 0.022 | 0.125 | 0.125 |
| | Upper CI | 57.143 | 3.099 | 3.099 | 0.188 | 0.475 | 0.475 |
| | t-Statistic | 69.250 | 124.144 | 124.144 | 41.706 | 91.966 | 91.966 |
| | P-Value | <0.001 | <0.001 | <0.001 | <0.001 | <0.001 | <0.001 |
| RLI2.3 | Observed | 0.000 | 1.000 | 1.890 | 0.500 | 0.468 | 0.125 |
| | Null Mean | 4.293 | 1.793 | 1.793 | 0.357 | 0.299 | 0.299 |
| | Lower CI | 0.000 | 1.000 | 1.000 | 0.222 | 0.120 | 0.120 |
| | Upper CI | 57.143 | 3.324 | 3.324 | 0.500 | 0.537 | 0.537 |
| | t-Statistic | 10.852 | 112.271 | 112.271 | 116.587 | 80.853 | 80.853 |
| | P-Value | <0.001 | <0.001 | <0.001 | <0.001 | <0.001 | <0.001 |
| RLI2.4 | Observed | 21.875 | 1.467 | 4.925 | 0.200 | 0.619 | 0.155 |
| | Null Mean | 14.620 | 2.978 | 2.978 | 0.240 | 0.409 | 0.409 |
| | Lower CI | 0.000 | 1.400 | 1.400 | 0.160 | 0.155 | 0.155 |
| | Upper CI | 56.250 | 4.925 | 4.925 | 0.320 | 0.697 | 0.697 |
| | t-Statistic | 33.312 | 86.235 | 86.235 | 164.817 | 75.085 | 75.085 |
| | P-Value | <0.001 | <0.001 | <0.001 | <0.001 | <0.001 | <0.001 |
| RLI3.1 | Observed | 0.000 | 1.000 | 3.007 | 0.430 | 0.523 | 0.218 |
| | Null Mean | 7.068 | 2.582 | 2.582 | 0.371 | 0.389 | 0.389 |
| | Lower CI | 0.000 | 1.000 | 1.000 | 0.231 | 0.217 | 0.217 |
| | Upper CI | 29.167 | 5.593 | 5.593 | 0.562 | 0.602 | 0.602 |
| | t-Statistic | 22.707 | 78.985 | 78.985 | 124.338 | 121.563 | 121.563 |
| | P-Value | <0.001 | <0.001 | <0.001 | <0.001 | <0.001 | <0.001 |

| | | | | | | | |
|---|---|---|---|---|---|---|---|
| RLI3.2 | Observed | 0.000 | 1.000 | 2.166 | 0.444 | 0.393 | 0.119 |
| | Null Mean | 12.214 | 1.793 | 1.793 | 0.315 | 0.296 | 0.296 |
| | Lower CI | 0.000 | 1.000 | 1.000 | 0.123 | 0.119 | 0.119 |
| | Upper CI | 57.143 | 3.442 | 3.442 | 0.494 | 0.534 | 0.534 |
| | t-Statistic | 21.196 | 109.053 | 109.053 | 86.350 | 82.328 | 82.328 |
| | P-Value | <0.001 | <0.001 | <0.001 | <0.001 | <0.001 | <0.001 |
| RLI3.3 | Observed | 0.000 | 1.305 | 1.667 | 0.451 | 0.461 | 0.258 |
| | Null Mean | 23.254 | 2.010 | 2.010 | 0.334 | 0.375 | 0.375 |
| | Lower CI | 0.000 | 1.222 | 1.222 | 0.174 | 0.245 | 0.245 |
| | Upper CI | 50.000 | 3.304 | 3.304 | 0.521 | 0.541 | 0.541 |
| | t-Statistic | 53.288 | 168.300 | 168.300 | 109.492 | 156.001 | 156.001 |
| | P-Value | <0.001 | <0.001 | <0.001 | <0.001 | <0.001 | <0.001 |
| RLI3.4 | Observed | 5.556 | 1.267 | 2.169 | 0.570 | 0.567 | 0.257 |
| | Null Mean | 6.869 | 2.000 | 2.000 | 0.469 | 0.399 | 0.399 |
| | Lower CI | 0.000 | 1.200 | 1.200 | 0.310 | 0.241 | 0.241 |
| | Upper CI | 22.222 | 3.524 | 3.524 | 0.570 | 0.589 | 0.589 |
| | t-Statistic | 28.036 | 122.675 | 122.675 | 210.183 | 124.028 | 124.028 |
| | P-Value | <0.001 | <0.001 | <0.001 | <0.001 | <0.001 | <0.001 |
| RLI4.1 | Observed | 11.905 | 1.246 | 5.293 | 0.249 | 0.472 | 0.298 |
| | Null Mean | 16.862 | 3.532 | 3.532 | 0.274 | 0.420 | 0.420 |
| | Lower CI | 0.000 | 1.246 | 1.246 | 0.194 | 0.295 | 0.295 |
| | Upper CI | 36.905 | 7.039 | 7.039 | 0.402 | 0.582 | 0.582 |
| | t-Statistic | 54.161 | 70.752 | 70.752 | 148.762 | 192.284 | 192.284 |
| | P-Value | <0.001 | <0.001 | <0.001 | <0.001 | <0.001 | <0.001 |
| RLI4.2 | Observed | 0.000 | 1.000 | 1.829 | 0.540 | 0.414 | 0.240 |
| | Null Mean | 13.185 | 1.958 | 1.958 | 0.397 | 0.364 | 0.364 |
| | Lower CI | 0.000 | 1.000 | 1.000 | 0.220 | 0.221 | 0.221 |
| | Upper CI | 46.154 | 3.597 | 3.597 | 0.580 | 0.519 | 0.519 |
| | t-Statistic | 30.915 | 123.150 | 123.150 | 125.655 | 156.177 | 156.177 |
| | P-Value | <0.001 | <0.001 | <0.001 | <0.001 | <0.001 | <0.001 |
| RLI4.4 | Observed | 0.000 | 1.400 | 1.902 | 0.640 | 0.435 | 0.382 |
| | Null Mean | 4.172 | 1.935 | 1.935 | 0.544 | 0.421 | 0.421 |
| | Lower CI | 0.000 | 1.400 | 1.400 | 0.430 | 0.363 | 0.363 |
| | Upper CI | 16.000 | 3.200 | 3.200 | 0.650 | 0.491 | 0.491 |
| | t-Statistic | 23.769 | 205.403 | 205.403 | 284.839 | 502.174 | 502.174 |
| | P-Value | <0.001 | <0.001 | <0.001 | <0.001 | <0.001 | <0.001 |
| VI1.1 | Observed | 0.000 | 1.000 | 1.667 | 0.375 | 0.314 | 0.126 |
| | Null Mean | 0.000 | 1.709 | 1.709 | 0.305 | 0.259 | 0.259 |
| | Lower CI | 0.000 | 1.000 | 1.000 | 0.250 | 0.121 | 0.121 |
| | Upper CI | 0.000 | 2.500 | 2.500 | 0.500 | 0.413 | 0.413 |
| | t-Statistic | NA | 147.756 | 147.756 | 120.416 | 104.247 | 104.247 |
| | P-Value | NA | <0.001 | <0.001 | <0.001 | <0.001 | <0.001 |
| VI1.3 | Observed | 11.111 | 1.638 | 3.007 | 0.248 | 0.481 | 0.375 |
| | Null Mean | 26.694 | 2.345 | 2.345 | 0.241 | 0.416 | 0.416 |
| | Lower CI | 0.000 | 1.659 | 1.659 | 0.157 | 0.333 | 0.333 |
| | Upper CI | 66.667 | 3.179 | 3.179 | 0.372 | 0.513 | 0.513 |
| | t-Statistic | 45.785 | 191.210 | 191.210 | 132.706 | 297.149 | 297.149 |
| | P-Value | <0.001 | <0.001 | <0.001 | <0.001 | <0.001 | <0.001 |

| | | | | | | | |
|---|---|---|---|---|---|---|---|
| VI1.4 | Observed | 0.000 | 1.333 | 3.000 | 0.417 | 0.423 | 0.233 |
| | Null Mean | 0.000 | 2.114 | 2.114 | 0.430 | 0.365 | 0.365 |
| | Lower CI | 0.000 | 1.333 | 1.333 | 0.327 | 0.243 | 0.243 |
| | Upper CI | 0.000 | 3.857 | 3.857 | 0.500 | 0.513 | 0.513 |
| | t-Statistic | NA | 118.396 | 118.396 | 234.073 | 161.563 | 161.563 |
| | P-Value | NA | <0.001 | <0.001 | <0.001 | <0.001 | <0.001 |
| VI2.1 | Observed | 16.667 | 1.267 | 3.797 | 0.380 | 0.544 | 0.261 |
| | Null Mean | 5.776 | 2.610 | 2.610 | 0.391 | 0.397 | 0.397 |
| | Lower CI | 0.000 | 1.200 | 1.200 | 0.250 | 0.230 | 0.230 |
| | Upper CI | 25.000 | 5.582 | 5.582 | 0.540 | 0.608 | 0.608 |
| | t-Statistic | 21.179 | 81.643 | 81.643 | 153.221 | 121.651 | 121.651 |
| | P-Value | <0.001 | <0.001 | <0.001 | <0.001 | <0.001 | <0.001 |
| VI2.2 | Observed | 0.000 | 1.000 | 1.679 | 0.497 | 0.424 | 0.319 |
| | Null Mean | 13.420 | 2.566 | 2.566 | 0.439 | 0.424 | 0.424 |
| | Lower CI | 0.000 | 1.308 | 1.308 | 0.314 | 0.345 | 0.345 |
| | Upper CI | 30.645 | 4.886 | 4.886 | 0.615 | 0.517 | 0.517 |
| | t-Statistic | 44.927 | 122.673 | 122.673 | 179.322 | 369.831 | 369.831 |
| | P-Value | <0.001 | <0.001 | <0.001 | <0.001 | <0.001 | <0.001 |
| VI2.3 | Observed | 0.000 | 1.000 | 1.503 | 0.500 | 0.338 | 0.216 |
| | Null Mean | 4.300 | 1.869 | 1.869 | 0.410 | 0.338 | 0.338 |
| | Lower CI | 0.000 | 1.000 | 1.000 | 0.333 | 0.223 | 0.223 |
| | Upper CI | 33.333 | 3.000 | 3.000 | 0.611 | 0.451 | 0.451 |
| | t-Statistic | 13.075 | 157.545 | 157.545 | 168.785 | 231.504 | 231.504 |
| | P-Value | <0.001 | <0.001 | <0.001 | <0.001 | <0.001 | <0.001 |
| VI2.4 | Observed | 0.000 | 1.800 | 1.400 | 0.480 | 0.274 | 0.404 |
| | Null Mean | 0.000 | 1.806 | 1.806 | 0.432 | 0.342 | 0.342 |
| | Lower CI | 0.000 | 1.400 | 1.400 | 0.333 | 0.257 | 0.257 |
| | Upper CI | 0.000 | 3.000 | 3.000 | 0.480 | 0.456 | 0.456 |
| | t-Statistic | NA | 203.195 | 203.195 | 270.625 | 227.467 | 227.467 |
| | P-Value | NA | <0.001 | <0.001 | <0.001 | <0.001 | <0.001 |
| VI3.3 | Observed | 0.000 | 1.000 | 1.000 | 0.750 | 0.281 | 0.281 |
| | Null Mean | 0.000 | 1.691 | 1.691 | 0.515 | 0.357 | 0.357 |
| | Lower CI | 0.000 | 1.400 | 1.400 | 0.367 | 0.307 | 0.307 |
| | Upper CI | 0.000 | 2.143 | 2.143 | 0.750 | 0.417 | 0.417 |
| | t-Statistic | NA | 243.866 | 243.866 | 187.419 | 532.929 | 532.929 |
| | P-Value | NA | <0.001 | <0.001 | <0.001 | <0.001 | <0.001 |
| VII1.1 | Observed | 0.000 | 1.000 | 1.563 | 0.521 | 0.405 | 0.225 |
| | Null Mean | 22.950 | 1.980 | 1.980 | 0.333 | 0.371 | 0.371 |
| | Lower CI | 0.000 | 1.000 | 1.000 | 0.166 | 0.228 | 0.228 |
| | Upper CI | 50.000 | 3.408 | 3.408 | 0.556 | 0.527 | 0.527 |
| | t-Statistic | 47.625 | 142.423 | 142.423 | 96.246 | 158.646 | 158.646 |
| | P-Value | <0.001 | <0.001 | <0.001 | <0.001 | <0.001 | <0.001 |
| VII1.2 | Observed | 0.000 | 1.000 | 3.298 | 0.508 | 0.558 | 0.224 |
| | Null Mean | 7.882 | 2.575 | 2.575 | 0.398 | 0.412 | 0.412 |
| | Lower CI | 0.000 | 1.000 | 1.000 | 0.203 | 0.216 | 0.216 |
| | Upper CI | 25.806 | 5.468 | 5.468 | 0.559 | 0.641 | 0.641 |
| | t-Statistic | 30.317 | 77.928 | 77.928 | 126.258 | 108.286 | 108.286 |
| | P-Value | <0.001 | <0.001 | <0.001 | <0.001 | <0.001 | <0.001 |

| | | | | | | | |
|---|---|---|---|---|---|---|---|
| VII1.3 | Observed | 0.000 | 1.000 | 2.118 | 0.165 | 0.377 | 0.123 |
| | Null Mean | 33.364 | 2.033 | 2.033 | 0.152 | 0.294 | 0.294 |
| | Lower CI | 0.000 | 1.000 | 1.000 | 0.066 | 0.122 | 0.122 |
| | Upper CI | 57.143 | 3.536 | 3.536 | 0.397 | 0.525 | 0.525 |
| | t-Statistic | 43.402 | 106.331 | 106.331 | 60.227 | 89.698 | 89.698 |
| | P-Value | <0.001 | <0.001 | <0.001 | <0.001 | <0.001 | <0.001 |
| VII1.4 | Observed | 0.000 | 1.000 | 2.534 | 0.180 | 0.396 | 0.126 |
| | Null Mean | 24.957 | 2.110 | 2.110 | 0.167 | 0.292 | 0.292 |
| | Lower CI | 0.000 | 1.000 | 1.000 | 0.080 | 0.120 | 0.120 |
| | Upper CI | 57.143 | 3.638 | 3.638 | 0.360 | 0.524 | 0.524 |
| | t-Statistic | 33.306 | 100.785 | 100.785 | 72.327 | 89.450 | 89.450 |
| | P-Value | <0.001 | <0.001 | <0.001 | <0.001 | <0.001 | <0.001 |
| VII2.1 | Observed | 0.000 | 1.445 | 1.445 | 0.500 | 0.281 | 0.296 |
| | Null Mean | 11.583 | 1.664 | 1.664 | 0.376 | 0.305 | 0.305 |
| | Lower CI | 0.000 | 1.333 | 1.333 | 0.222 | 0.271 | 0.271 |
| | Upper CI | 66.667 | 2.219 | 2.219 | 0.500 | 0.359 | 0.359 |
| | t-Statistic | 22.145 | 252.570 | 252.570 | 138.421 | 472.712 | 472.712 |
| | P-Value | <0.001 | <0.001 | <0.001 | <0.001 | <0.001 | <0.001 |
| VII2.2 | Observed | 0.000 | 1.000 | 4.914 | 0.180 | 0.469 | 0.129 |
| | Null Mean | 5.618 | 3.218 | 3.218 | 0.208 | 0.322 | 0.322 |
| | Lower CI | 0.000 | 1.000 | 1.000 | 0.140 | 0.116 | 0.116 |
| | Upper CI | 27.273 | 6.052 | 6.052 | 0.420 | 0.643 | 0.643 |
| | t-Statistic | 16.433 | 67.018 | 67.018 | 83.798 | 74.684 | 74.684 |
| | P-Value | <0.001 | <0.001 | <0.001 | <0.001 | <0.001 | <0.001 |
| VII2.3 | Observed | 25.000 | 1.490 | 2.667 | 0.277 | 0.384 | 0.423 |
| | Null Mean | 25.905 | 2.302 | 2.302 | 0.302 | 0.413 | 0.413 |
| | Lower CI | 0.000 | 1.496 | 1.496 | 0.194 | 0.368 | 0.368 |
| | Upper CI | 50.000 | 3.592 | 3.592 | 0.488 | 0.467 | 0.467 |
| | t-Statistic | 68.406 | 169.677 | 169.677 | 127.663 | 737.489 | 737.489 |
| | P-Value | <0.001 | <0.001 | <0.001 | <0.001 | <0.001 | <0.001 |
| VII3.1 | Observed | 16.667 | 1.573 | 1.342 | 0.340 | 0.332 | 0.328 |
| | Null Mean | 22.488 | 2.041 | 2.041 | 0.357 | 0.370 | 0.370 |
| | Lower CI | 0.000 | 1.267 | 1.267 | 0.220 | 0.320 | 0.320 |
| | Upper CI | 50.000 | 2.810 | 2.810 | 0.560 | 0.425 | 0.425 |
| | t-Statistic | 46.322 | 207.662 | 207.662 | 119.503 | 607.874 | 607.874 |
| | P-Value | <0.001 | <0.001 | <0.001 | <0.001 | <0.001 | <0.001 |
| VII3.2 | Observed | 0.000 | 1.333 | 1.828 | 0.604 | 0.432 | 0.385 |
| | Null Mean | 7.333 | 1.905 | 1.905 | 0.535 | 0.421 | 0.421 |
| | Lower CI | 0.000 | 1.389 | 1.389 | 0.382 | 0.364 | 0.364 |
| | Upper CI | 22.000 | 3.000 | 3.000 | 0.653 | 0.491 | 0.491 |
| | t-Statistic | 34.557 | 197.595 | 197.595 | 226.553 | 507.742 | 507.742 |
| | P-Value | <0.001 | <0.001 | <0.001 | <0.001 | <0.001 | <0.001 |
| VII3.3 | Observed | 0.000 | 1.627 | 1.000 | 0.625 | 0.226 | 0.447 |
| | Null Mean | 6.019 | 1.861 | 1.861 | 0.457 | 0.365 | 0.365 |
| | Lower CI | 0.000 | 1.000 | 1.000 | 0.297 | 0.219 | 0.219 |
| | Upper CI | 30.769 | 3.085 | 3.085 | 0.656 | 0.529 | 0.529 |
| | t-Statistic | 19.767 | 130.349 | 130.349 | 155.375 | 148.882 | 148.882 |
| | P-Value | <0.001 | <0.001 | <0.001 | <0.001 | <0.001 | <0.001 |

| | | | | | | | |
|---|---|---|---|---|---|---|---|
| VII4.1 | Observed | 0.000 | 1.000 | 1.000 | 0.406 | 0.222 | 0.222 |
| | Null Mean | 35.483 | 1.825 | 1.825 | 0.267 | 0.318 | 0.318 |
| | Lower CI | 0.000 | 1.250 | 1.250 | 0.125 | 0.264 | 0.264 |
| | Upper CI | 66.667 | 2.312 | 2.312 | 0.500 | 0.367 | 0.367 |
| | t-Statistic | 47.248 | 231.301 | 231.301 | 82.354 | 449.865 | 449.865 |
| | P-Value | <0.001 | <0.001 | <0.001 | <0.001 | <0.001 | <0.001 |
| VII4.2 | Observed | 19.231 | 1.232 | 1.764 | 0.521 | 0.471 | 0.260 |
| | Null Mean | 17.396 | 1.870 | 1.870 | 0.393 | 0.376 | 0.376 |
| | Lower CI | 0.000 | 1.154 | 1.154 | 0.213 | 0.247 | 0.247 |
| | Upper CI | 46.154 | 2.998 | 2.998 | 0.544 | 0.535 | 0.535 |
| | t-Statistic | 44.195 | 162.698 | 162.698 | 135.778 | 153.993 | 153.993 |
| | P-Value | <0.001 | <0.001 | <0.001 | <0.001 | <0.001 | <0.001 |

**Table S4** Non-significant Generalized Additive Mixed Models (GAMMs) testing the relationship between network descriptors and environmental variables, for 144 trophic networks. Response variables are shown for each model fitted with smoothing functions: elevation, date, time of the day and maximum microclimatic temperature for June and July (Tmax_06 and Tmax_07). The established random (re) factors include Site-Habitat and Transect (4 nested transects for each habitat).

| Response variable | Predictor | Edf (smooth) | P-value | | Adjusted $r^2$ | Deviance explained | Family | Link function |
|---|---|---|---|---|---|---|---|---|
| Generality HL | Intercept | - | <0.001 | *** | 0.153 | 27.7% | quasi-Poisson | log |
| | Elevation | 1.206 | 0.91545 | n. s | | | | |
| | Tmax_07 | 1.000 | 0.69112 | n. s | | | | |
| | Tmax_06 | 1.000 | 0.39874 | n. s | | | | |
| | Date | 1.000 | 0.99299 | n. s | | | | |
| | Time | 1.441 | 0.65326 | n. s | | | | |
| | Transect [re] | 5.552 | 0.29775 | n. s | | | | |
| | Site [re] | 3.484 | 0.00785 | ** | | | | |
| Vulnerability LL | Intercept | - | <0.001 | *** | 0.267 | 39.4% | quasi-Poisson | log |
| | Elevation | 1.000 | 0.1158 | n. s | | | | |
| | Tmax_07 | 1.000 | 0.3184 | n. s | | | | |
| | Tmax_06 | 1.000 | 0.8947 | n. s | | | | |
| | Date | 1.000 | 0.2042 | n. s | | | | |
| | Time | 1.441 | 0.1091 | n. s | | | | |
| | Transect [re] | 8.564 | 0.1001 | n. s | | | | |
| | Site [re] | 3.520 | 0.0129 | * | | | | |
| Robustness LL | Intercept | - | <0.001 | *** | 0.225 | 38.1% | Gaussian | Identity |
| | Elevation | 1.559 | 0.5213 | n. s | | | | |
| | Tmax_07 | 1.000 | 0.5132 | n. s | | | | |
| | Tmax_06 | 1.277 | 0.8748 | n. s | | | | |
| | Date | 1.000 | 0.7820 | n. s | | | | |
| | Time | 1.000 | 0.7299 | n. s | | | | |
| | Transect [re] | 11.593 | 0.0372 | * | | | | |
| | Site [re] | 1.633 | 0.1763 | n. s | | | | |
| Modularity Q | Intercept | - | <0.001 | *** | 0.265 | 40.9% | Gaussian | Identity |
| | Elevation | 1.201 | 0.4850 | n. s | | | | |
| | Tmax_07 | 1.000 | 0.4399 | n. s | | | | |
| | Tmax_06 | 1.000 | 0.4511 | n. s | | | | |
| | Date | 1.000 | 0.2976 | n. s | | | | |
| | Time | 1.142 | 0.5124 | n. s | | | | |
| | Transect [re] | 11.85 | 0.0129 | * | | | | |
| | Site [re] | <0.001 | 0.5644 | n. s | | | | |
